# Optimal balancing of clinical factors in large scale clinical RNA-Seq studies

**DOI:** 10.1101/2021.06.30.450639

**Authors:** Austin W.T. Chiang, Vahid H. Gazestani, Mia G. Altieri, Benjamin P. Kellman, Srinivsa Nalabolu, Tiziano Pramparo, Karen Pierce, Eric Courchesne, Nathan E. Lewis

## Abstract

Omics technologies are ubiquitous in biomedical research. However, improper sample selection is an often-overlooked complication with large omics studies, resulting in confounding effects that can disrupt the internal validity of a study and lead to false conclusions. Here, we present a method called ***BalanceIT***, which uses a genetic algorithm to identify an optimal set of samples with balanced clinical factors for large-scale omics experiments. We apply our approach to two large RNA-Seq studies in autism (1) to find a post-hoc balanced sample set among an imbalanced study, and (2) to design an optimal study that allows for efficient batch correction. Our approach leads to near-perfect estimates of differential gene expression, superior performance of pathway-level enrichment analysis, and consistent network dysregulation patterns of autism symptom severity. These results provide empirical support for the importance of balanced experimental design, and ***BalanceIT*** will be invaluable for large-scale study design and batch effect correction.

## Introduction

High throughput technologies enable scientists to develop large-scale studies with molecular-scale insights into complex biological systems involved in diverse areas of biological and clinical research.^1–4^ In clinical research, large-scale studies have a tremendous impact on clinical development and practice.^5^ In particular, RNA-Seq and other omics technologies^6–8^ have enabled more comprehensive investigation of disease mechanisms, and have paved the way for translating exciting biomedical research into clinical practice.^9–11^ To enable this, it is crucial that experimental and analytical practices accurately quantify gene expression levels.^12–15^ Although many techniques (e.g., quality control^16^, normalization^17–19^, replicate control^20,21^, and spike-in control^22,23^) have addressed challenges leading to erroneous quantification of transcript abundance, potential sources of error remain in RNA-Seq library design and analysis. For example, many large clinical studies intend to assay hundreds or thousands of samples, prepared in batches or individual “plates” (e.g., 96-well plates to facilitate sample handling). Since each sample may be associated with dozens of clinically relevant patient characteristics, it is important to design the study to avoid confounding these with batch assignment. However, it remains unclear to what extent study design^24–26^ in large RNA-Seq experiments impacts results of the gene expression analysis.

One largely unexplored direction is the consideration of clinical factor balancing in sample selection and sequencing library design. In clinical trials, the randomized controlled trial (RCT) is recognized as important to establish validity of causal associations between the intervention and studied subjects.^27–29^ RCTs are usually performed by randomly allocating study subjects into different treatment groups,^30^ in which each subject is enrolled into a group with an equal probability. The advantage of using randomization is the possibility of relaxing the underlying assumption that the majority of subject characteristics are not biased between groups under study and provide a basis for inference.^31–33^ In large-scale RNA-Seq studies, wherein samples are allocated in multiple batches, experimental design often overlooks the balances between samples. While samples may be randomly allocated to different batches or plates for sequencing experiments, the assumption of balanced subject characteristics can be violated when a bias in subject characteristic occurs between groups.^34–36^ While the probability that any one characteristic is unbalanced during randomization is small, the large number of clinical features greatly increases the chance that some features will be unbalanced in a study and ultimately correlate erroneously with diagnosis, treatment, or important measured features. Thus, the biased characteristic generates unwanted variation in gene expression and confounds biological variation leading to the batch effect problem^37,38^. This idea, however, has not been directly studied in regard to how experimental design impacts global gene expression and downstream analysis and the resulting biological conclusions.^39–42^

Here we propose an experimental balancing strategy for large omics experiments, ***BalanceIT***, that employs a genetic algorithm (GA)^43,44^ to identify an optimal set of samples with balanced clinical factors. We adopt a concept deviating drastically from that of the conventional sample-based randomization. In essence, the novelty of the concept is that the scoring function is entirely a covariate-based optimization approach. Thus, library balancing will be achieved by searching for an optimal set of samples with balanced covariate factors. This reduces the chance that global gene expression is influenced by confounding factors from sample organization and covariates of patient characteristics. Most importantly, we show that conventional randomization may not be reliable enough for large clinical datasets. Here we also evaluate the behavior of different library design strategies in a large-scale autism RNA-Seq dataset (1,566 subjects). We first used our method to design a balanced dataset, from which we sequenced 549 subjects to assess the performance of techniques for removing batch effects and performing downstream analyses (e.g., differential expression, enrichment analysis, and network activity analysis). Second, we show ***BalanceIT*** can be used for post-hoc dataset correction on an older, unbalanced dataset of 695 ASD subjects to show how one can identify balanced subsets. We further demonstrate that ***BalanceIT*** is not just superior to conventional randomization in designing a balanced library for RNA-Seq experiment, but also improves upon state-of-the-art batch effect correction methods in accurately inferring gene expression levels; thus, it will be invaluable for identifying causal associations from large scale omics studies.

## Results

### Sample randomization is not enough to eliminate confounding factors in large scale clinical studies

To introduce the library design problem, we examined simulated RNA-Seq data (see **Methods**). We consider two different library designs (**Text S1**). The first, a balanced library, intends to assay gene expression of 200 subjects where the different diagnosis groups (case and control) are randomly allocated to different plates, and three different clinical covariates were randomly assigned to subjects in a balanced way (**Figure 1A, i**). Specifically, the clinical variates and plate assignments are independent from the diagnosis groups. Note that, the clinical covariates are the confounding factors but often overlooked in the study design. On the contrary, plates are usually human designated in the study design process. The second, an imbalanced library, intends to assay gene expression of 300 subjects where the different diagnosis groups are allocated to different clinical variates or plates in a biased way (**Figure 1B, i**). To achieve this, we added 100 subjects to the balanced library (200 subjects), in which the newly added subjects have their clinical variates or plates associated with the diagnosis groups (**Methods**). The resulting library thus shows correlations between clinically related patient characteristics or plates and the diagnosis groups.

**Figure 1.**
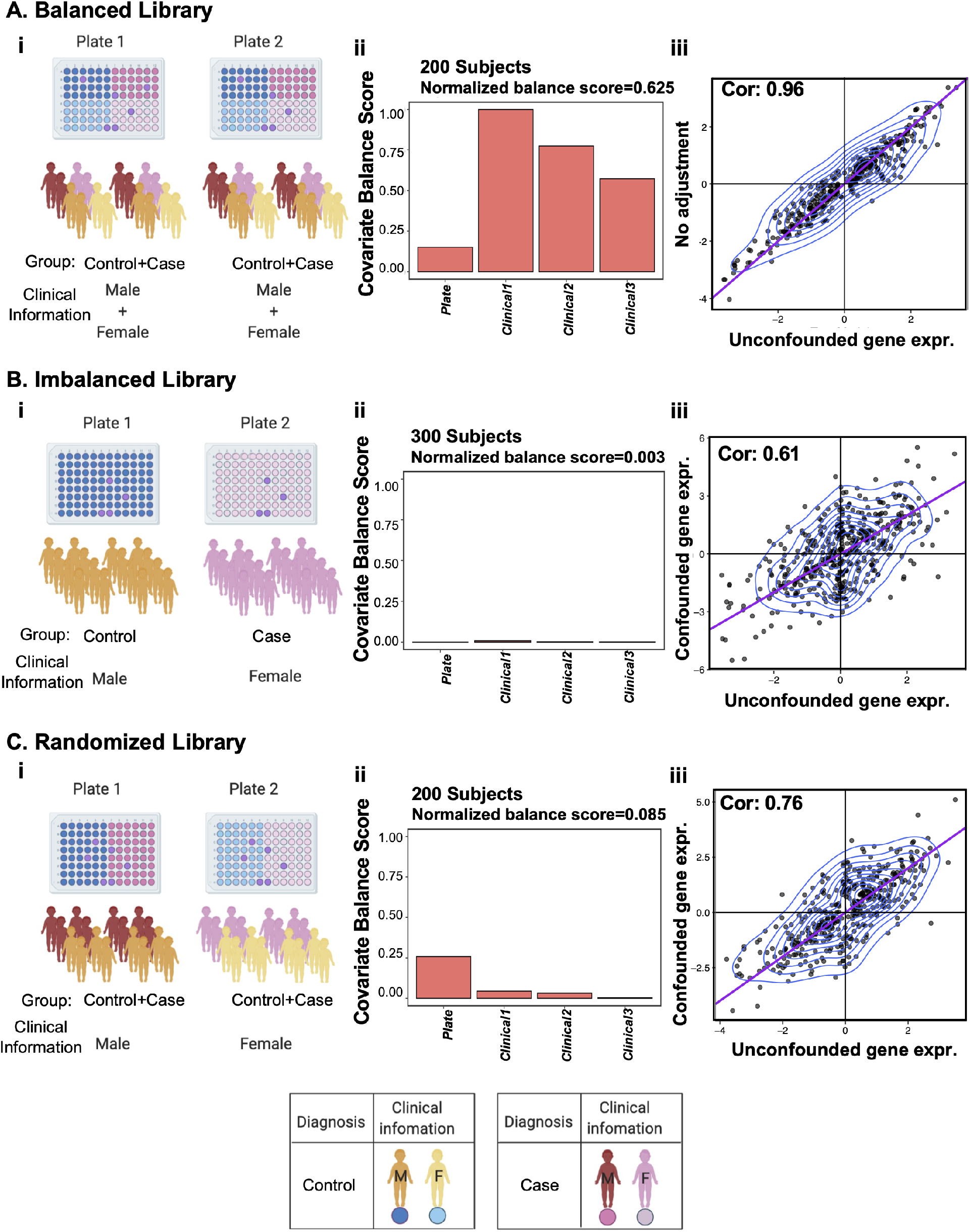
Effects of sequencing library design. Impact of study design on gene expression is examined for three different library designs: balanced library (**A**), imbalanced library (**B**), and randomized library (**C**). (**i**) A schematic of the RNA-Seq library design. (**ii**) Balance score of batch variables. (**iii**) Fold change pattern of differentially expressed genes between the unconfounded simulated gene expression and batch-effect-confounded gene expression.

The balancing score of each factor (i.e., clinical covariate or plate) is assessed by testing statistical distributions of this factor in two different diagnosis groups, and the overall balance score of the study design is the normalized total covariate balance scores (**Text S1**). Our results show that the overall balancing for the balanced library is high (normalized balance score=0.625; **Figure 1A-ii**), but the overall balancing for the imbalanced library is very low (normalized balance score=0.003) (**Figure S1B-ii)**. We further simulated expression for 1,000 genes for both library designs. The first 400 genes were designed to be differentially expressed between diagnosis groups, and the first 500 genes were simulated to be affected by clinical variates or plates (**Methods**). To investigate the effect of the study design strategy on gene expression, we compared the fold change patterns of DE genes between the unconfounded simulated gene expression and batch-effect-confounded gene expression from different library designs (**Text S1)**. Our results show that similar fold change patterns were observed in the balanced library design with a Pearson’s correlation coefficient (r) of 0.96 (**Figure 1A, iii**), but the fold change patterns were much more affected in the imbalanced library design (r=0.61) (**Figure 1B, iii**). Moreover, the batch effect correction methods perform worse than the gene expression corrected by known batches when quantifying the concordance of DEG rank (**Figure S1**). As expected, these results indicate that imbalanced library designs substantially impact global gene expression, as unwanted variation cannot be easily removed, leading to inaccurate gene expression levels.

Intriguingly, we investigated whether standard randomization of sample organization and sequencing libraries is effective in balancing patient characteristics in a large-scale clinical study. We utilized the above simulated imbalanced library (300 subjects) as the pool of candidate samples to test different balancing approaches. Then, we randomly selected the same number of samples for the two different diagnosis groups (i.e., 100 case and 100 control subjects) and randomly allocated them to different sequencing plates (**Figure 1C-i**). Our results show that randomization can ensure the plates are independent from the diagnosis groups. However, the other clinical covariates cannot be ensured to be independent from diagnosis groups, and the overall balancing for this randomized library remains low (normalized balance score=0.085) (**Figure 1C-ii**). Although the gene expression might be accurately corrected with all known batches (clinical 1-3) (r = 0.99), the fold change pattern of the confounded gene expression has deviated substantially from the real gene expression (r=0.76) (**Figure 1C-iii**). Unfortunately, if one does not know all the batches but just controls the plates, the gene expression might not be accurately corrected by batch effect correction methods (r ≥ 0.86) (**Figure S3C**). These results demonstrate that randomization library might fail in managing a balanced library for large-scale clinical study with many covariates.

### Accuracy increases with removal of samples that introduce imbalance

Next, we tested if sample organization and sequencing libraries can be designed to balance patient characteristics in a large-scale clinical study. Thus, we designed the library using our GA-based method-***BalanceIT***, to identify an optimal set of samples with all the clinical factors balanced across all sequencing library plates (**Methods**). To allow comparison to the randomized library, we applied ***BalanceIT*** to select subjects from the same imbalanced library (**Figure 2A**). We found the resulting library (174 subjects) has all the clinical covariates and plates independent from the diagnosis groups. However, the resulting library contains fewer subjects than the original designed 200 subjects in the balanced library since we made the trade-off between library size and library balancing. Specifically, the overall balancing for the ***BalanceIT*** library now achieves the maximum balance (normalized balance score=1.00; **Figure 2B**) compared to the balance of the original balanced library (normalized balance score=0.625; **Figure 1A**). Examining the gene expression pattern, the fold change pattern shows great consistency with the real gene expression for both corrected and without batch corrected analyses (r=0.98) (**Figure 2C**). Moreover, even if one does not know all clinical variates (clinical 1-3) but just controls the plates, the gene expression still can be accurately corrected by all batch effect correction methods we tested (r ≥ 0.98) (**Figure S3D**). Furthermore, the ***BalanceIT*** library presents more accurate and stable of DEG ranking (**Figure S2B**). Moreover, our results showed that the balanced library outperforms the imbalanced library, and the fully balanced ***BalanceIT*** library has better performance than the randomization library in the downstream RNA-Seq analysis (**Figure S4C**). A better-balanced library design would yield more accurate enriched gene sets, while a less balanced library design may lead to erroneous conclusions from incorrect enriched gene sets **(Text S2)**. All these results suggest that unwanted variation is well managed and removed through balancing the RNA-Seq library.

**Figure 2.**
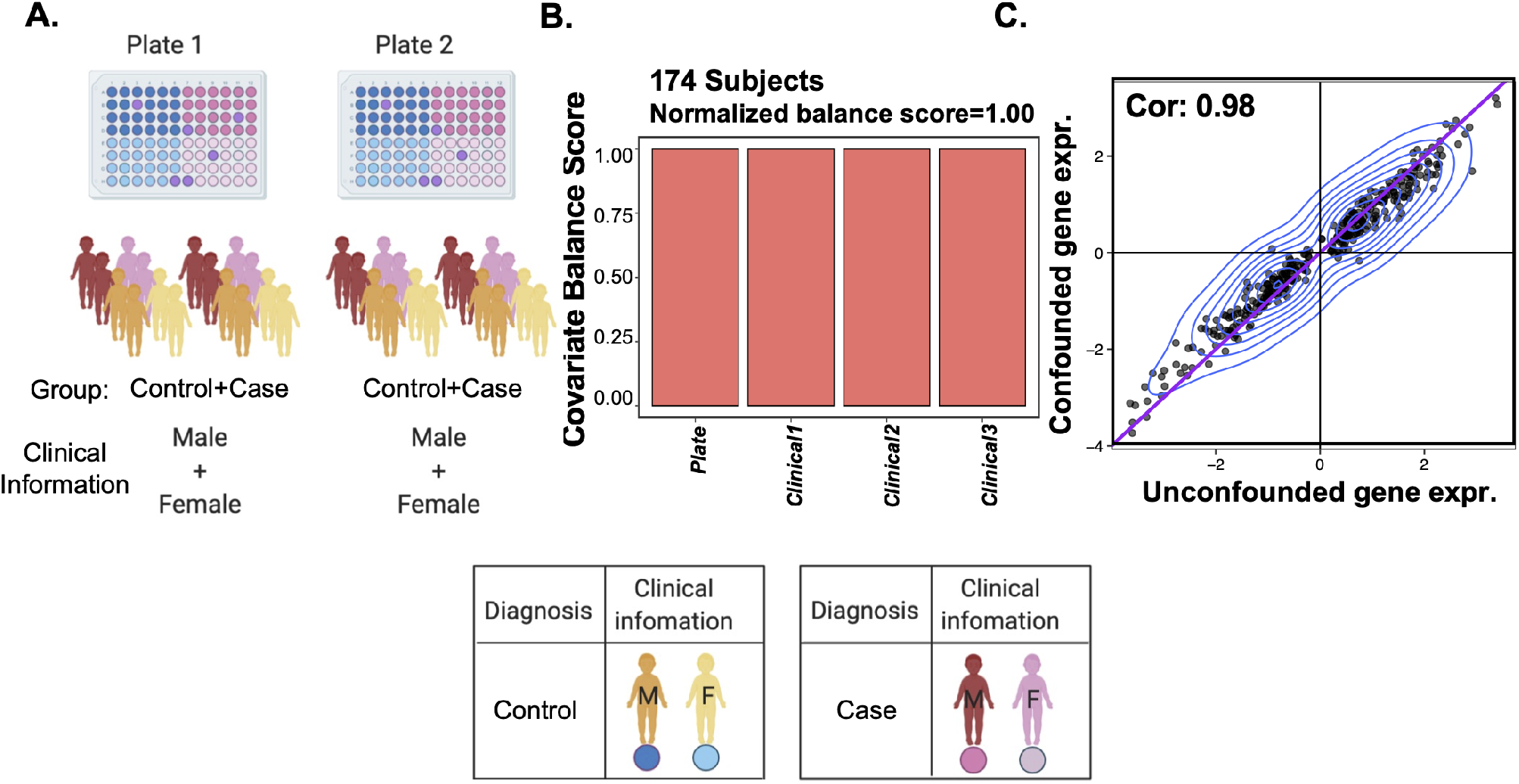
Effects of sequencing library design using BalanceIT. Impact of library design on gene expression is examined for the library designs using BalanceIT. (**A**) A schematic of the RNA-Seq library design. (**B**) Balance score of batch variables. (**C**) Fold change pattern of differentially expressed genes between the unconfounded simulated gene expression and batch-effect-confounded gene expression. All the notations are the same as in **Figure 1**.

### Datasets with imbalances can be corrected with *BalanceIT*

We evaluated the performance of sequencing library design with two large-scale clinical datasets for studying transcriptional changes in ASD. The first dataset, referred to here as *IBL*, assayed 678 samples, but exhibits an imbalanced library design (subjects were substantially biased on gender and diagnosis across different sequencing library plates). The second dataset, *ACE*, included 1,780 samples (1,308 subjects) in 6 different diagnostic groups: autism spectrum disorder (ASD), typically developing (TD), language developmental delay (LD), developmental delay (DD), ASD feature subjects (ASDfeat), and the other subjects (Others)). Importantly, these two large-scale clinical datasets were handled the same prior to the sequencing library design, and include 23 clinical characteristics, all measured at the UCSD Autism Center of Excellence (**Methods**). Thus, we used these datasets to study our library balancing strategy.

It is worth noting that, while we intended to sequence 1,780 RNA samples, our aim is to control batch effects that might plague the gene expression dataset. Therefore, we employed BalanceIT to create a balanced, non-batch contaminated experiment with the 1,780 samples before any of these samples were sequenced. Here, we sequenced 549 of the 1,780 samples, in which the 549 samples with balanced covariates between different diagnosis groups. These 549 newly sequenced samples allowed us for investigating how batch effect might impact on the performance of differential gene expression, pathway-level enrichment analysis, and network dysregulation patterns of autism symptom severity.

To fair compare the performance, we compared the 549 sequenced samples to results from the *IBL* dataset (678 samples). We found that the overall balancing for the *IBL* library has low balance (normalized balance score=0.166; **Figure 3A**). Randomization test showed that the balance of this library can be optimized (p>0.99; **Figure S6**). To study if we can correct the gene expression of the *IBL* data, we used ***BalanceIT*** to select a set of samples with balanced covariates between different diagnosis groups. Our results shows that the overall balancing for the optimized *IBL* library substantial increase (3.17-fold) its balance (normalized balance score=0.528; **Figure 3B**).

**Figure 3.**
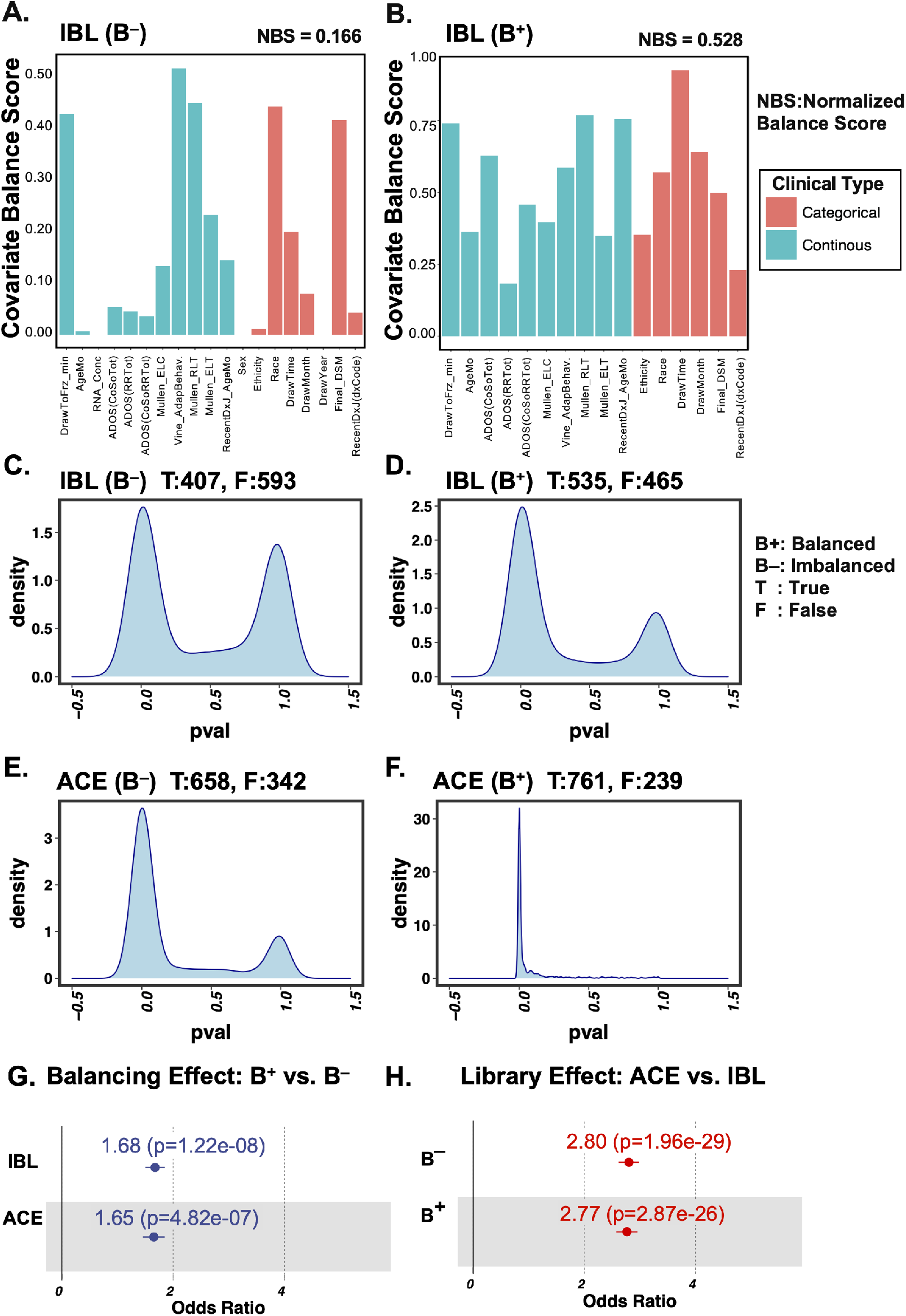
Gene co-expression analysis in the ASD-specfic (DE-ASD) network. Four different strategies for RNA-Seq data analysis: The balancing results for all covariates of the IBL data **(A)** and the **BalanceIT** balanced IBL data **(B)**. **C**) imbalanced library design (IBL) without balancing correction, **D**) IBL with balancing correction, **E**) balanced library design without balancing correction, and**F**) balanced library design with balancing correction. **G**) Odds ratio comparing IBL(B^+^) vs. IBL(B^-^) and ACE(B^+^) vs. ACE(B^-^) about their gene co-expression magnitudes. **H**) Odds ratio comparing ACE (B^-^) vs. IBL (B^-^) and ACE (B^+^) vs. IBL (B^+^) about their gene co-expression magnitudes. ‘**B**’: the balancing correction; ‘T’: the number of randomly sampled ASDprofiles with significantly (Wilcoxon test: p<0.05) higher gene coexpression than the randomly sampled TDs; ‘F’: the number of randomly sampled ASD profiles does not have significantly (p≥0.05) higher gene co-expression than the randomly sampled TDs. The randomization testing was conducted by randomly sampling 50 ASDs and 50 TDs from a total of 231 samples (**Table S1**) for 1,000 times.

Interestingly, our previous study demonstrated that DEGs in leukocytes from ASD subjects, measured via microarrays, could be used to build a molecular interaction network (DE-ASD network) that showed gene co-expression that increased with ASD severity.^45^ In this study, we wondered if the increased co-expression activity seen in ASD subjects could be replicated in the larger RNA-Seq datasets described in this current study, and if the signal is sensitive to subject balancing. First, we tested the performance of the gene co-expression analysis in the DE-ASD network using the *IBL* dataset. Specifically, we took the 231 overlapping samples between the *IBL* dataset (678 samples) and the ACE balanced dataset (549 samples), and we tested if the transcriptomes show increasing gene co-expression from toddlers with ASD by two different strategies (**Figure 3, C and D**) of RNA-Seq data analysis: 1) *IBL*(B^−^)–imbalanced library design without balance correction and 2) *IBL*(B^+^)–imbalanced library design with balance correction. We found that the balance corrected data significantly improved the signal of gene co-expression in the DE-ASD network for the *IBL* dataset (odds ratio=1.68) (**Figure 3G**). Thus, our results demonstrated that imbalanced datasets can be corrected with our ***BalanceIT***.

### High quality datasets can be designed with optimal balancing

Next, we applied ***BalanceIT*** to create a fully balanced sequencing library from our ASD cohort, selected from Dataset II–*ACE* data. Specifically, we used ***BalanceIT*** to optimize the library with balanced covariates between different diagnosis groups. ***BalanceIT*** identified 789 subjects with well-balanced covariates distributed between groups: ASD vs. TD (normalized balance score=0.636), TD vs. LD (normalized balance score=0.713), TD vs. DD (normalized balance score=0.715), and TD vs. ASDfeat (normalized balance score=0.679) (**Figures S7**). Importantly, since we aim to identify an optimal solution that the samples have all their covariates well-balanced across the RNA-Seq plates, we further used ***BalanceIT*** to successfully allocate the identified 789 subjects onto 13 balanced plates (normalized balance score=0.971; **Figure 4A**). Additionally, since magnetic resonance imaging (MRI) has been known as a powerful tool for investigating brain structural changes in children with ASD, we further used ***BalanceIT*** to allocate 358 subjects onto 6 balanced plates that are enriched with MRI data for RNA-Seq (normalized balance score=0.694; **Figure 4B**). Randomization test shows that the overall balance of the optimized library is statistically significant with p value=6.6e-03 (**Figure S8**). Examining individual covariates, we found each covariate shows good balance scores across the 6 plates (**Figures 4B, S9 and S10**). It is worth noting that the ‘covariate balance score’ is assessed by testing statistical distributions of this factor in two different diagnosis groups. While some scores seem not high (covariate balance score=0.2, 0.3, … etc.), the distributions of these covariates are statistically the same (p>0.05). For example, the continuous covariates of DrawToFrz_min, ADOS RRTot, and ADOS CoSo (covariate balance score=0.278, 0.370 and 0.202, respectively; **Figure 4C**), and categorical covariates-Sex, Ethnicity, and Extraction Date (covariate balance score=0.592, 0.629, and 0.640, respectively; **Figure 4D**). Thus, ***BalanceIT*** has successfully balances the RNA-Seq library to allows for further control of batch effects in large-scale RNA-Seq experiments (**Figure 4**).

**Figure 4.**
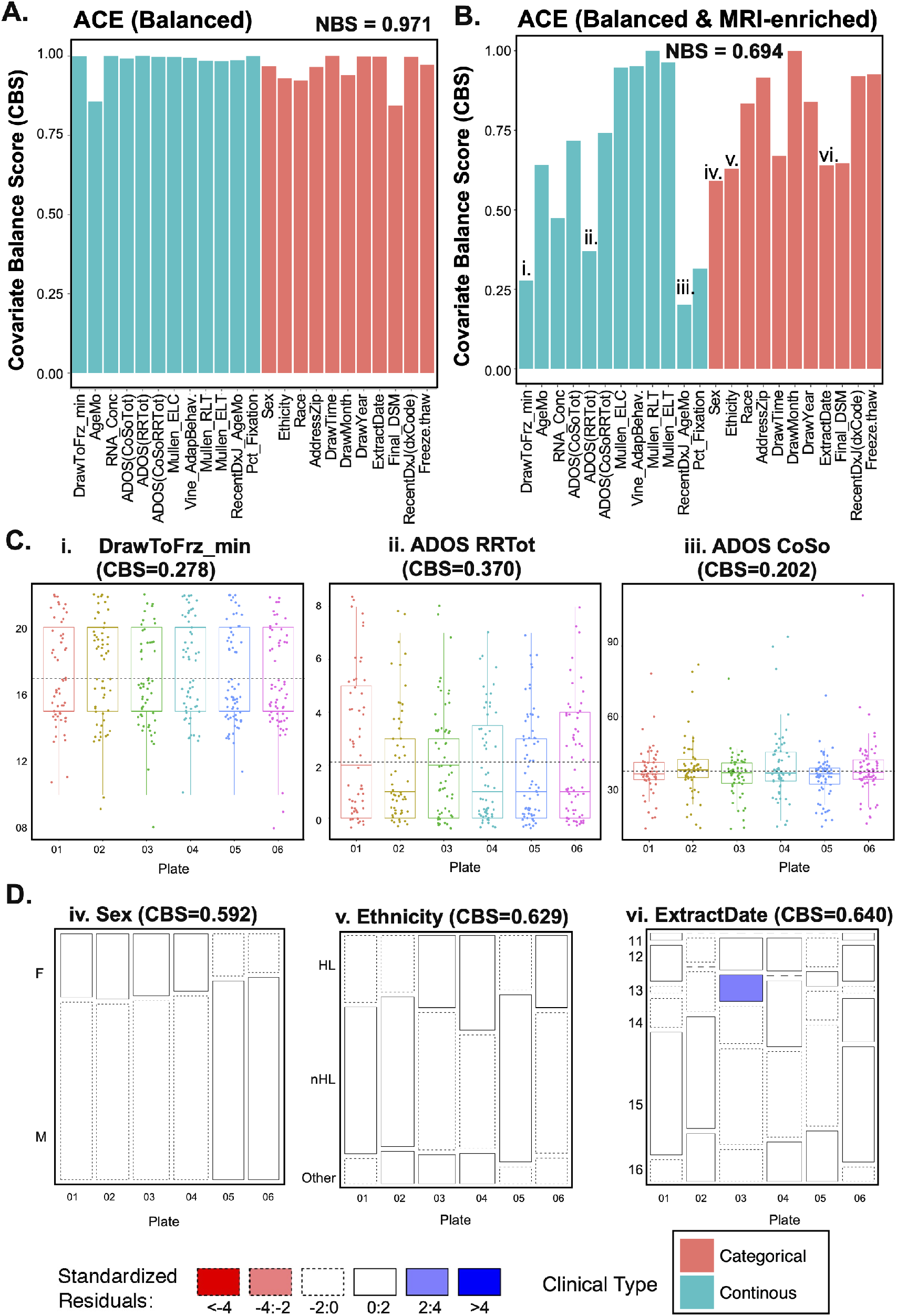
BalanceIT successfully balances covariates across 13 RNA-Seq library plates for samples from the ACE dataset. **A)** The balancing results for all categorical and continuous covariates. The y-axis represents the balancing score of each covariate. **B**) The balancing results for the selected continuous covariates-Age (months), RNA-Concentration and ADOS CoSo. **C**) The balancing results for the selected categorical covariates-Sex, Ethnicity, and Race. NBS: normalized balance score.

To demonstrate the power of ***BalanceIT*** in RNA-Seq data analysis, we first validate on differential expression patterns for the IBL and ACE-MRI datasets (**Text S3**). Our results show that ACE-MRI can reproduce consistent differential expression patterns of two public datasets-tData^46^ and Natneur253^45^ (R=0.76 and R=0.79), but the *IBL* design gene expression cannot be accurately corrected by the batch effect correction methods (R=0.46 and R=0.26) (**Figures S13-14**). These analyses corroborated the effectiveness of balanced library design on batch effect control in clinical RNA-Seq data. Next, we tested the performance of the aforementioned gene co-expression analysis in the DE-ASD network using the *ACE* and *IBL* datasets. For a fair comparison, we used the same 231 overlapping samples described above, and tested if the transcriptomes show increasing DE-ASD network gene co-expression in ASD subjects, using four different strategies (**Figure 3, C and D**) of RNA-Seq data analysis: 1) *ACE*(B^-^)–balanced library design without batch effect correction and 2) *ACE*(B^+^)–balanced library design with batch effect correction. Interestingly, **Figures 3E-F** show that: 1) the batch effect correction significantly improved the signal of gene co-expression in the DE-ASD network for the *ACE* datasets (odds ratio=1.65); 2) the balanced library design played a dominant role in enhancing the signal of DE-ASD network gene coexpression on both datasets (odds ratio>2.75); and 3) the maximum improvement in the gene coexpression signal is observed with the balanced library design with batch effect correction compared with the imbalanced library design without batch effect correction (odds ratio=4.64, p=3.87e-59). Thus, balancing the dataset during experimental design allows the effective removal of batch effects in large RNA-Seq experiments.

In addition to increased gene co-expression in the DE-ASD network, Gazestani et al.^45^ further found the gene co-expression activity further increased with increasing symptom severity among ASD subjects. We further tested if the balanced library design impacts the reported correlation of gene coexpression magnitude and ASD severity, as measured by the Autism Diagnostic Observation Schedule social affect (ADOS-SA) severity score for the *IBL* dataset and the balanced *ACE* dataset with batch effect correction. The *IBL* dataset shows a poor prediction of ASD symptom severity (**Figure 5, A and B**); however, the balanced *ACE* dataset showed better association of ASD social severity with the DE-ASD network dysregulation (**Figure 5, C and D**). The findings using the balanced *ACE* dataset are consistent with the results of the original study^45^, despite it having been originally done using microarrays and on a different set of subjects. Our results suggest that an imbalanced library design is confounded by experimental setup factors correlating with clinical covariates and diagnosis; therefore, it will be challenging to separate the experimental factors from the clinical factors in the *IBL* dataset.

**Figure 5.**
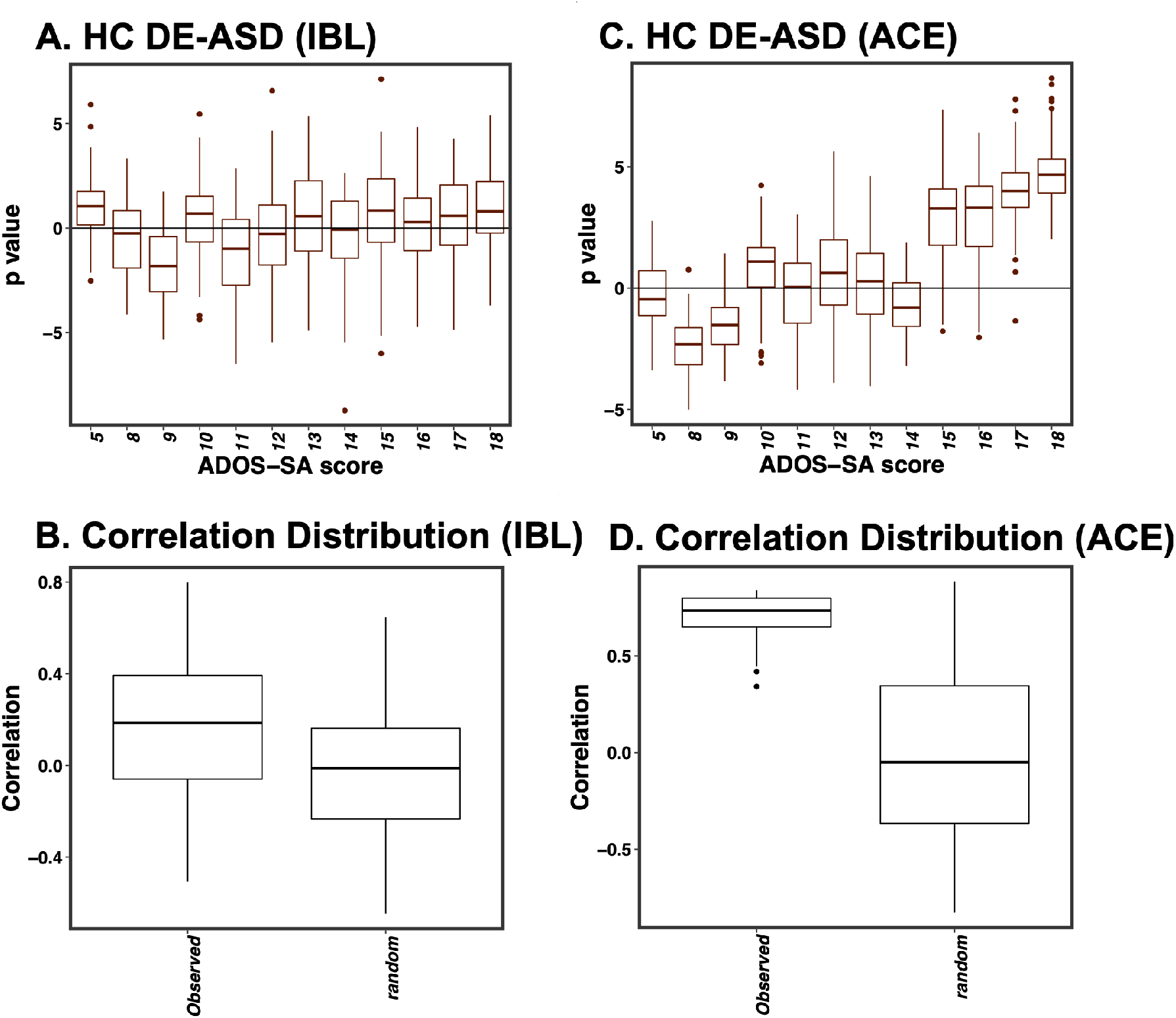
Co-expression magnitude of the DE-ASD network correlates with ASD symptom severity. The relative coexpression magnitude of the DE-ASD networks compared with TD networks using the imbalanced library design of IBL dataset (i and iii) and the balanced library design of ACE dataset (ii and iv). The relative activity level was estimated by comparing the co-expression strength of interactions in the DE-ASD network between toddlers with ASD and with TD. For each severity group, n = 20 ASD samples in that ADOS-SA range were randomly selected and compared with n = 20 random TD samples, iterating 1,000 times. Significance of the trend was evaluated by 10,000 permutations of the ADOS-SA scores in toddlers with ASD (two-sided P < 10^-6^; permutation test). Boxplots represent the median (horizontal line), lower and upper quartile values (box), the range of values (whiskers) and outliers (dots).

## Discussion and Conclusions

Managing unwanted variation and eliminating batch effects is foundationally critical for valid biological conclusions.^39–42^ Here we sought to optimize sample selection for balanced designs for managing confounding effects stemming from sample organization and covariates of sample characteristics. We demonstrated that imbalanced library designs substantially impact the quantification of global gene expression, resulting in erroneous RNA-Seq analysis results and falsely reporting clinically relevant differences. Unfortunately, unwanted variations from confounding factors are difficult to remove from imbalanced library designs, and this can lead to false positive or negative conclusions about gene expression levels. Here, we introduced a new optimization-based method, ***BalanceIT***, to identify an optimal set of samples with balanced covariate factors and organize these for easy removal of batch effects in large studies. Our method outperforms conventional randomization, suggesting that there is merit to systematic library balancing approaches.

As a proof-of-principle, a GA was employed for the optimization that exhaustively searches for balanced solutions. GAs are commonly used to generate high-quality solutions in optimization and search research.^43,44^ Even though GA has the undeniable merit of offering valuable efficiency into the optimization process, its limitations include their demand on computational resources required for converging to the final optimal balanced solution. However, this limitation can be alleviated by integrating with advanced sampling approaches such as Latin hypercube sampling^47^ and multiple ellipsoid-based sampling^48^ that have been developed to efficiently search for parameter solutions from high dimensional spaces. Replacing our time-consuming simulation step with these efficient parameter-searching methods, one can collect great amounts of patient characteristics (parameters) from large clinical datasets, and then apply ***BalanceIT*** on those datasets to find the optimal solution of balanced samples.

Our approach could be particularly useful in clinical studies with a large number of patient characteristics that can have complicated confounding impact on gene expression. The results of simulated gene expression experiments show that an optimized library design provides a significantly better mean for accurately inferring gene expression levels and improving downstream functional analyses (e.g., differential expression analysis and gene set enrichment analysis) than randomization. We also performed ***BalanceIT*** on the ASD datasets and demonstrated proper balancing was needed to reproduce a previously reported correlation between transcriptional dysregulation with increased ASD symptom severity^45^. The results are generally supportive of both simulated gene expression datasets and real clinical datasets. They indicate that major source of unwanted variation can be easily eliminated using the library design identified by ***BalanceIT***. These findings suggest that ***BalanceIT*** enables researchers to better control batch effects and to draw valid biological conclusions in large scale clinical studies.

## Acknowledgments

This work was supported by National Institute of Mental Health (NIMH) R01-MH110558 (E.C., N.E.L.) and generous funding from the Novo Nordisk Foundation through the Technical University of Denmark (NNF20SA0066621; N.E.L.).

## Online Methods

### Simulated dataset

In this study, we simulated gene expression datasets for four different scenarios of library designs:

- *Scenario I (Balanced Library)*–this dataset composed of 200 subjects with three different clinical variables and two different disease groups. We simulated 1,000 genes on 2 plates for these 200 subjects, in which the gene expression follows a negative binomial model for count-based RNA-sequencing data^49^. The model parameters were estimated from the Zebrafish data as described in Leek et al.^50^. The RNA-sequencing count data were simulated with the Polyester Rpackage^51^. The first 400 genes were designed to be differentially expressed between diagnosis groups, and the first 500 genes were simulated to be affected by clinical covariates or plates. All the group, plate, and batch variables (clinical variables) were not correlated in this scenario (**Figure 1A**).
- *Scenario II (Imbalanced Library)*–we add 100 subjects to *Dataset I*. The added 100 subjects have strong correlation between plate and the disease group, and moderate correlations between clinical variables and the disease group (**Figure 1B**). The count-based RNA-sequencing data was further simulated as described above.
- *Scenario III (Randomized Library)*–we randomly select 200 subjects from *Dataset II* and randomly assign them into two different plates. These 200 subjects therefore did not show correlation between plate and the disease group, but they show correlations between clinical variables and the disease group (**Figure 1C**). The count-based RNA-sequencing data was simulated as described above.
- *Scenario IV (BalanceIT Library)*–we applied ***BalanceIT*** to *Dataset II* and identify an optimal set of 174 subjects with balanced clinical variables on two plates (**Figure 2B**). The count-based RNA-sequencing data was further simulated as described above.

### Data processing and differential gene expression analysis of the simulated gene expression datasets

The count-based RNA-sequencing data was processed by the following procedures. The limma package^52^ was first applied on quantile-normalized count data for differential expression analysis in which moderated t-statistics were calculated by robust empirical Bayes methods^53^. We estimated batch effects using six different batch effect correction methods: 1) supervised svaseq, 2) unsupervised svaseq, 3) principal components analysis, 4) RUV with control probes, 5) known batch correction (true simulated batch variables), and 6) no adjustment. Sequencing plate was used as a categorical covariate and corrected using one of the above batch effect correction method. Genes with Benjamini-Hochberg corrected *P* <0.05 were deemed as DEGs. For investigating the effect of the library design strategy on gene expression, we compared the fold change patterns of DE genes between real gene expression and confounded gene expressions from different library designs.

### *BalanceIT* — A genetic algorithm-based library balancing method

The GA-based approach is developed to search the optimal design of RNA-Seq library with each experimental factor is balanced. **Figure S15** illustrates the flowchart of the proposed GA-based method optimization framework. (A) ***Initial population***. The initial population is filled with 150 individuals, in which each individual (*N*) consists of *l* chromosomes {*P*_1_,*P*_2_,…,*P_l_*}, each with a subset of samples (i.e., genes) from the large RNA-Seq dataset {*S*_1_,*S*_2_,…,*S_m_*} annotated with known experimental factors {*F*_1_,*F*_2_,…,*F_n_*}. (B) ***Evaluation***. To assess whether the distributions of experimental factors in the *l* sets of samples of individual (*N_i_*) differ, each experimental factor is evaluated by performing the ANOVA (Analysis of Variance) test for continuous factor and by the Chi-square (*χ*^2^) test for categorical factor of the *l* sets {*P*_1_,*P*_2_,…, *P_l_*} of samples in *N_i_*. The *p*-value of statistical test is considered as the ‘balance’ score of the testing experimental factor, in which ‘1’ denotes balanced (the two distributions are identical) and ‘0’ means not balanced (the two distributions are different). The fitness score is defined as the summation of all the balance scores in *N_i_*, which is used to be maximized in the GA optimization process. (C) ***Stop criteria***. The stopping criterion is set to a preset maximum of 100 generations. (D) ***Selection***. The population is sorted by descending fitness values, and the top 30% population (45 individuals) are considered as candidates to select two individuals as parents (*N_i_* and *N_j_*) for further crossover and mutation to generate a new individual of next-generation population. This step iterative 150 times to generate the same population size as the initial population. (E) ***Crossover***. A single crossover point on both parents’ chromosome strings {*P*_1_,*P*_2_} is selected, and all data beyond that point in either chromosome strings is swapped between the two parent organisms. The resulting chromosome strings are the children (*N’*). (F) ***Mutation***. Mutation used to maintain genetic diversity by swapping one point gene values in the two children’s (*N’*) chromosomes.

### Data processing and differential gene expression analysis of the ASD gene expression datasets

In this study, we performed transcriptomics analysis of two datasets:

- *Dataset I*–the IBL dataset has imbalanced library design (subjects were substantially biased on genders across different plates) on a total of 678 subjects. The imbalanced library was further used for RNA sequencing experiment.
- *Dataset II*–the *ACE* dataset from Autism Center of Excellence-UCSD, contains 1,308 subjects in 6 different autistic diagnosis groups (autism spectrum disorder (ASD), typical developing (TD), language developmental delay (LD), developmental delay (DD), ASD feature subjects (ASDfeat), and the other subjects (Others)). We performed ***BalanceIT*** to identify 789 subjects (balanced *ACE* library) with well-balanced covariates distributed between groups. From the balanced *ACE* library (789 unique toddlers) together with 266 longitudinal samples and 248 technical replicated samples (**Figures S11**), we sequenced 549 subjects (6 plates) to assess the performance of techniques for removing batch effects and performing downstream analyses.

Quality of RNA-Seq samples was examined using FastQC^54^. RNA-Seq data were mapped and quantified using STAR^55^ and HTSeq^56^, respectively. The limma package^52^ was then applied on quantile-normalized count data for differential expression analysis in which moderated t-statistics were calculated by robust empirical Bayes methods^53^. Batch (sequencing plate) was used as a categorical covariate and corrected using Combat^57^ batch effect correction tool. Genes with Benjamini-Hochberg corrected *P* <0.05 were deemed as DEGs, which were used for further analyses (e.g., fold change comparison, network activity analysis, and gender effect analysis). Pearson’s correlation coefficient was used for the comparison of fold changes across datasets.

For validating differential expression patterns of the above two datasets, we performed transcriptomics analysis of two published datasets:

- *Validation Dataset I*–the *tData* was reported by Pramparo et al.^46^, which was composed of 128 subjects (n= 84 ASD and 44 TD subjects).
- *Validation Dataset II*–the *NatNeur253* was reported by Gazestani et al.^45^, which was composed of 253 subjects (n= 84 ASD and 44 TD subjects).

To ensure that results are not affected by variations in the processing procedures, we performed our preprocessing analyses (e.g., quality control and normalization) exactly the same as those reported by Gazestani et al.^45^. The limma package^52^ was then applied on the quantile-normalized data for differential expression analysis in which moderated t-statistics were calculated by robust empirical Bayes methods^53^. Sample batch was used as a categorical covariate and corrected using Combat^57^ batch effect correction tool. Genes with Benjamini-Hochberg corrected *P* <0.05 were deemed as DE.

### Gene set enrichment analysis (GSEA) and enrichment strength analysis

GSEA was performed using the Broad Institute GSEA software^58^. A ranked list of genes (adjusted p-values < 0.05) was made using the differential expression values (Fold change in the log2 scale) from differential gene expression analysis were run through the GSEA pre-ranked protocol. GSEA-pre-rank analysis was processed to detect significant molecular signature terms (‘Hallmark’ (50) and ‘KEGG’ (186) gene sets from the MSigDB were used here) for the differential expressed genes (p value < 0.05). Note that, the criteria for considering a molecular signature term as significant are: 1) after FDR false discovery correction, molecular signature terms with p-value < 0.05 and FDR adjusted p-values less than 0.1; and 2) there are >10 genes presented in our gene list of this molecular signature terms.

### Network Activity Analysis

To further compare the results of the balancing, the network signature of the data was produced to see how ASD network interactions change based on the data. The ASD network is represented as pairs of genes, signifying an interaction in the network. To generate the network signature the counts of the RNA-Seq data were first normalized using VST, which computes a function that when applied to the data the variability of the function is not related to the mean value of the data. These counts are then separated by sample according to their diagnosis, resulting in two normalized sets of samples, one for ASD samples and another for TD samples, ASD = x and TD = y. To determine the strength of the interactions in the ASD network, correlations between the pairs were commuted using Pearson’s correlation for both sets of samples. Finally, the absolute values of the ASD interactions and TD interactions were subtracted from each other, to show how the ASD network interactions are impacted by diagnosis.

### Constructing the male and female consensus networks and associate modules with clinical traits

A consensus network^59^ represents a single network arising from multiple sources of data constructed from the weighted average of correlation matrices from both the male and female in this study. Followed the protocols of weighted gene co-expression network analysis (WGCNA)^60^, we created consensus networks for male and female. Co-expression networks were constructed with the Combat corrected genes in the *ACE* dataset (n= 197 ASD and 192 TD subjects). Specifically, Pearson’s correlation coefficients were first calculated for all the genes in the dataset and the correlation matrix of the entire gene dataset was obtained. Genes were further hierarchically clustered, and modules were determined by using a dynamic tree-cutting algorithm^61^. The consensus network and module generation were performed in WGCNA with the following changes to the default settings for consensus network generation: β = 7, deepSplit = 1, cutHeight = 0.25, and a minimum module size of 30 genes. Module identifiers in the male network were then changed to match the most similar module in the female network based on gene overlap. The gene expression profile for a consensus module was summarized by a single representative gene (named, ‘eigengene’ (E)), i.e., a module could be characterized by a single representative gene.^59^ To identify the modules that are strongly correlated with the clinical traits, we further constructed the module-trait relationships by calculating Pearson’s correlations between the module signature (eigengene expression) and the clinical traits.^62^ The gene significance was calculated as the absolute value of the association between the expression profile and each clinical trait. The module membership was defined as the correlation between the expression profile and each module signature.

### Differential expressed network module analysis

To identify the modules that show differential expressed between male and females, we defined the differentially expressed modules by three criteria: 1-2) the correlation between eigengene_i_ and trait_j_, Cor(E_i_, T_j_), should be greater than 0.2 and statistical significance with p value less than 0.05 for at least one network module (either male or female network module), and 3) the fold change of the correlation between eigengene and trait in the comparison (Cor(E_i_, T_j_)^(male)^/Cor(E_i_, T_j_)^(female)^) between male and female modules should be greater/less than 50%.

