## Supplementary Information for "Optimal balancing of clinical factors in large scale clinical RNA-Seq studies"

### Supplementary Texts


**Text S1.** Unwanted variation in large-scale clinical datasets.
**Text S2.** Library balancing impacts RNA-Seq analysis.
**Text S3.** Validation on differential expression patterns for the IBL and ACE-MRI datasets


### Supplementary Table


**Table S1.** Overview of the 231 samples shared between the IBL dataset and the ACE-MRI RNA-Seq dataset.


### Supplementary Figures


**Figure S1.** Effects of balanced and imbalanced library design on simulated gene expression.
**Figure S2.** Effects of randomized and BalanceIT library design on simulated gene expression.
**Figure S3.** Fold change pattern of differentially expressed genes for different batch effect correction methods.
**Figure S4.** Impact of different library designs on post-sequencing analysis.
**Figure S5.** P-value distribution for differentia expression test between diagnosis groups (ASD vs. TD).
**Figure S6.** Randomization test of the IBL library design for the ASD dataset with 678 samples.
**Figure S7.** Balancing results of the BalanceIT library design for the ACE-MRI dataset with selected 789 subjects.
**Figure S8.** Randomization test of the ACE-MRI library design for the ASD dataset with 358 samples.
**Figure S9.** Balancing results for the continuous covariates of the ASD dataset.
**Figure S10.** Balancing results for the continuous covariates of the ASD dataset.
**Figure S11.** Flow chart of the ACE balanced library in the RNA sequencing experiment.
**Figure S12.** Reproducibility analysis of the observed DE pattens for *NatNeur253* and *tData*.
**Figure S13.** Reproducibility analysis of the observed DE pattens for *IBL* and *ACE* datasets.
**Figure S14.** Reproducibility analysis of the observed DE pattens.
**Figure S15.** Overview of the Genetic Algorithm Optimization framework of *BalanceIT*.

**Text S1. Unwanted variation in large-scale clinical datasets**

For the simulated gene expression datasets, we consider two different library designs (**Figure S1**). The first, a balanced library, intends to assay gene expression of 200 subjects where the different diagnosis groups (case and control) are allocated to different plates, and clinical covariates were randomly assigned to subjects in a balanced way (**Figure S1A, i-iii**). Note that, the clinical covariates are the confounding factors but often overlooked in the study design. On the other hand, plates are usually human designated in the study design process. Therefore, we defined the often overlooked 'clinical covariates' as 'batches' throughout the study. Specifically, the batches or plates are independent from the diagnosis groups. The balancing score of each factor (i.e., clinical covariate or plate) is assessed by testing statistical distributions of this factor in two different diagnosis groups, and the overall balance score of the study design is the normalized total covariate balance scores. The overall balancing for this library is high (normalized balance score=0.625). The second, an imbalanced library, intends to assay gene expression of 300 subjects where the different diagnosis groups are allocated to different batches or plates in a biased way (**Figure S1B, i-iii**). To achieve this, we added 100 subjects to the balanced library (200 subjects), in which the newly added subjects have their batches or plates associated with the diagnosis groups (Materials and Methods). The resulting library thus shows correlations between clinically related patient characteristics or plates and the diagnosis groups, and the overall balancing for this library is very low (normalized balance score=0.003).

For both library designs, we simulated expression for 1,000 genes (**Methods**). The first 400 genes were designed to be differentially expressed between diagnosis groups, and the first 500 genes were simulated to be affected by batches or plates. To investigate the effect of the library design strategy on gene expression, we compared the fold change patterns of DE genes between the unconfounded simulated real gene expression and batch-effect-confounded gene expression from different library designs. As expected, similar fold change patterns were observed in the balanced library design with a Pearson's correlation coefficient (r) of 0.96 (**Figure S1A, v**). If the confounded gene expression data were further corrected by known batches, the fold change patterns are almost the same (r= 0.99; **Figure S1A, vi**). Even without knowledge of all batches (clinical 1-3) but just plate controls, the gene expression still can be accurately corrected by several well-known batch effect correction methods ($r \geq 0.95$) (**Figure S3B**). However, examining the concordance of the top 400 DEGs, we found that for each DEG ranking, supervised and unsupervised svaseq outperformed two other common methods: principal component analysis (PCA), and remove unwanted variation (RUV) with empirical controls, and not adjusted for batch effects (**Figure S1A, iv**). Surprisingly, the fold change patterns were much more affected in the imbalanced library design (r=0.61) (**Figure S1B, v**). The accurate gene expression can be called only if the confounded gene expression were corrected by known batches (r= 0.98; **Figure S1B, vi**). Unfortunately, if one does not know all batches (clinical 1-3) but just controls for plates, the gene expression cannot be accurately corrected by established batch effect correction methods ($r \leq 0.77$) (**Figure S3A**). Indeed, the batch effect correction methods perform worse than the gene expression corrected by known batches when quantifying the concordance of DEG rank (**Figure S1B, iv**). These results indicate that imbalanced library designs substantially impact global gene expression, as unwanted variation cannot be easily removed, leading to inaccurate gene expression levels.

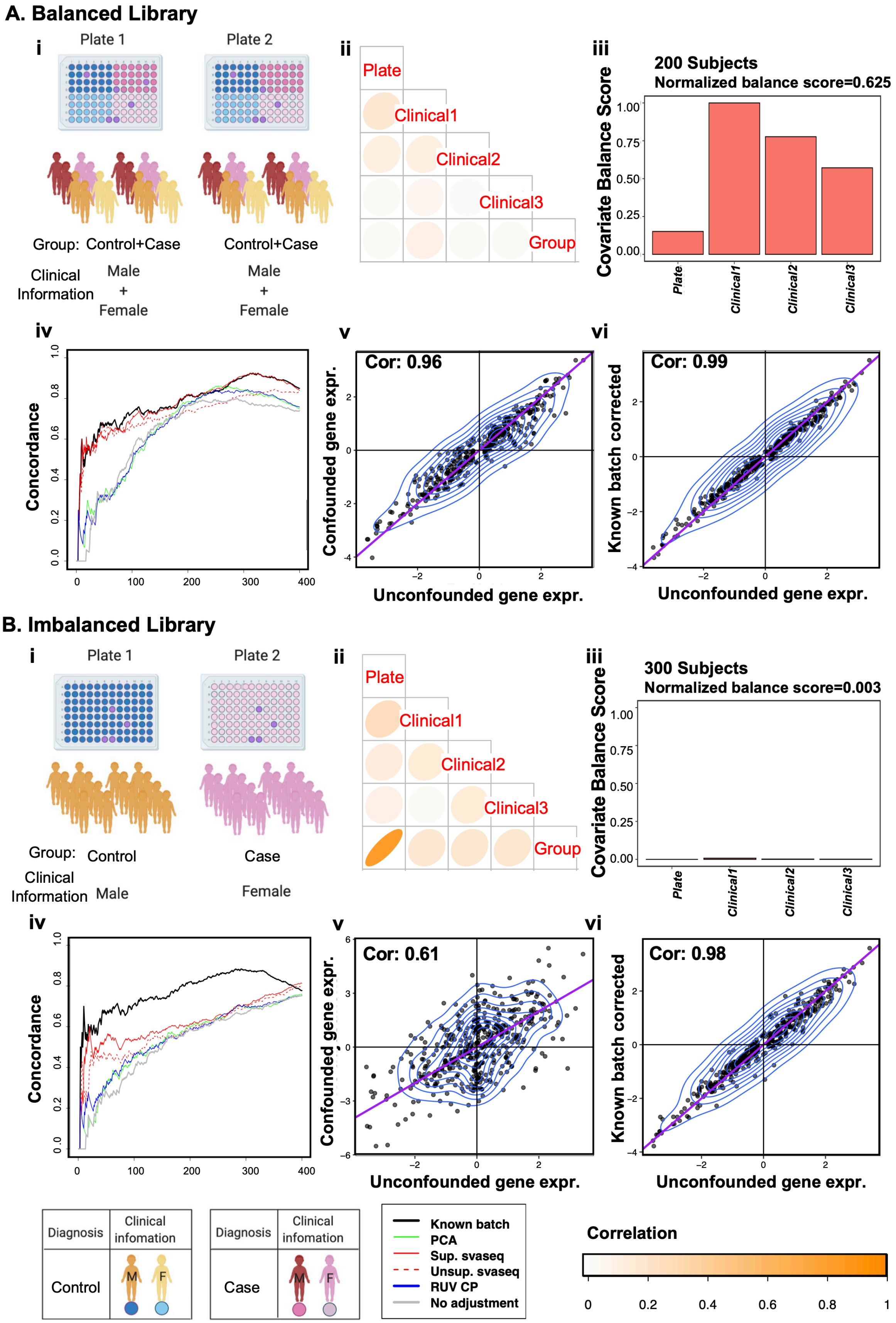


***Figure S1. Effects of balanced and imbalanced library design on simulated gene expression.*** *We quantify the impact of sample organization in library design on simulated gene expression for two different library designs: balanced library*

*(**A**) and imbalanced library (**B**). (**i**) The diagram illustrates the clinical information and their diagnostic group for subjects in the RNA-Seq library design. (**ii**) Correlation between simulated batch variables (plate and clinical information–clinical 1-3) and diagnostic group. (**iii**) Balance score, which quantifies whether each covariate is balanced in the two diagnostic groups– ASD and TD (see Materials and Methods). Proximity to '1' denotes a more balanced distribution, while '0' means completely imbalanced. (**iv**) The concordance plot shows the fraction of differentially expressed genes that are concordant between gold standard (unconfounded gene expression) and different batch effect correction methods (known batch, principal component analysis (PCA), supervised svaseq, unsupervised svaseq, remove unwanted variation (RUV) using empirical control probes, and no adjustment for the confounded gene expression). (**v**) The density plot shows the fold change pattern of differentially expressed genes between real gene expression and batch-confounded gene expression of the library design. (**vi**) The density plot shows the fold change pattern of differentially expressed genes between real gene expression and known batch corrected gene expression.*

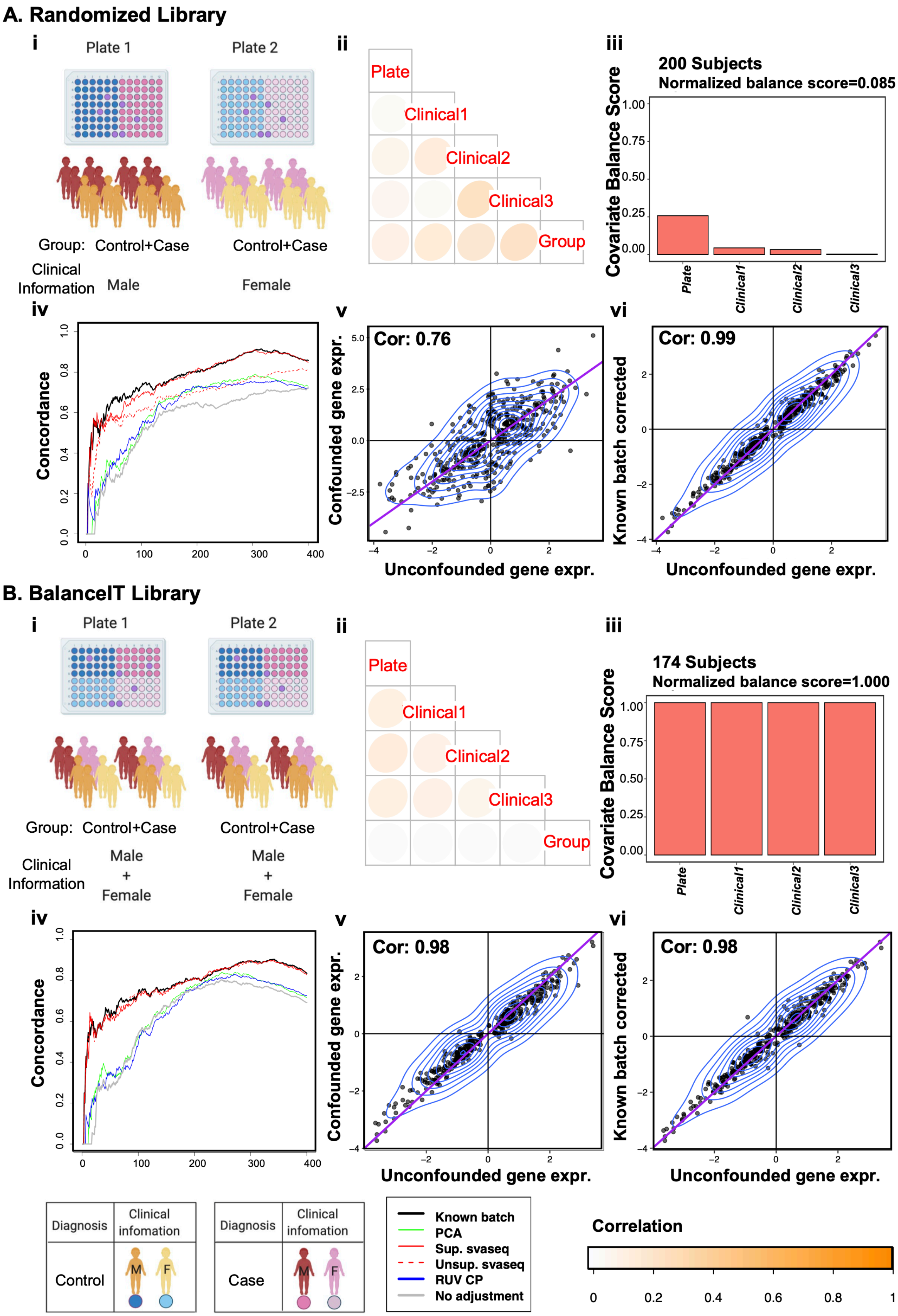


***Figure S2. Effects of randomized and BalanceIT library design on simulated gene expression.*** *We quantify the impact of sample organization in library design on simulated gene expression for two different library designs: balanced library*

*(**A**) and imbalanced library (**B**). (**i**) The diagram illustrates the clinical information and their diagnostic group for subjects in the RNA-Seq library design. (**ii**) Correlation between simulated batch variables (plate and clinical information–clinical 1-3) and diagnostic group. (**iii**) Balance score, which quantifies whether each covariate is balanced in the two diagnostic groups– ASD and TD (see Materials and Methods). Proximity to '1' denotes a more balanced distribution, while '0' means completely imbalanced. (**iv**) The concordance plot shows the fraction of differentially expressed genes that are concordant between gold standard (real gene expression) and different batch effect correction methods (known batch, principal component analysis (PCA), supervised svaseq, unsupervised svaseq, remove unwanted variation (RUV) using empirical control probes, and no adjustment for the confounded gene expression). (**v**) The density plot shows the fold change pattern of differentially expressed genes between real gene expression and batch-confounded gene expression of the library design. (**vi**) The density plot shows the fold change pattern of differentially expressed genes between real gene expression and known batch corrected gene expression.*

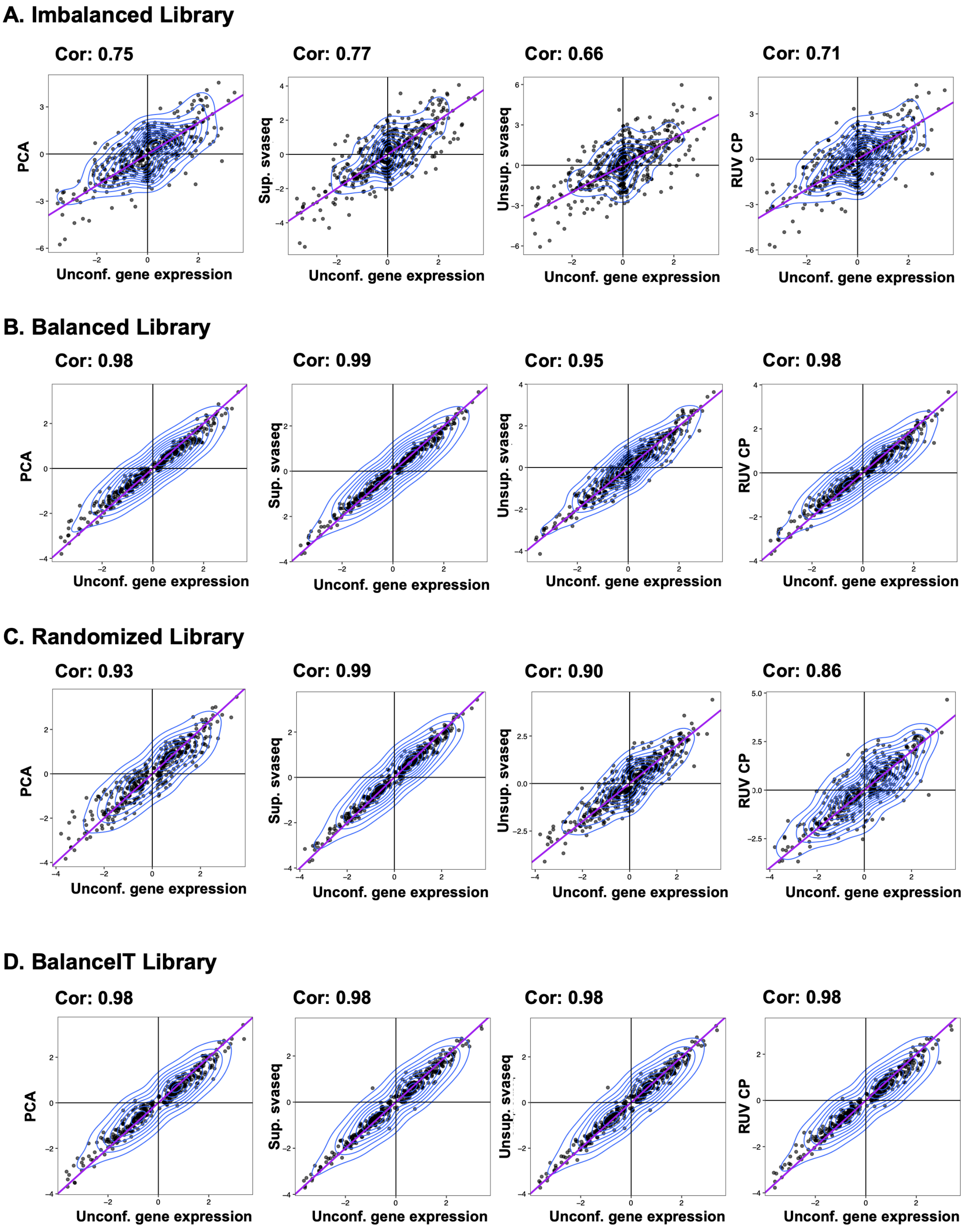


***Figure S3. Fold change pattern of differentially expressed genes for different batch effect correction methods.*** *Four different RNA-Seq library designs:* ***A****) imbalanced library design,* ***B****) balanced library design,* ***C****) randomized library design, and* ***D****) BalanceIT library design. For each library design, the confounded gene expression were corrected by four batch effect correction methods: principal component analysis (PCA), supervised svaseq, unsupervised svaseq, remove unwanted variation (RUV) using control probes. 'Unconf. gene expression' denotes unconfounded gene expression.*

### Text S2. Library balancing impacts RNA-Seq analysis

Two important applications of RNA-Seq are the analysis of differential gene expression (DEG) and gene set enrichment analyses (GSEA) for comparing different clinical groups (e.g., ASD vs. TD). To study the impact of sequencing library design on post-sequencing analysis, we compared the DEGs and enriched gene sets. Here we defined DEGs using two criteria: (1) less stringent, with FDR adjusted p-values <0.05 (**Figure S4A)**, and (2) more stringent, considering both FDR adjusted p-values <0.05 and an absolute fold change >1.5 (**Figure S4B)**. We found the balanced library design outperforms the imbalanced library in terms of sensitivity, specificity, and accuracy. The excellent performance of the balanced library stems from fewer falsely positive DEGs. Interestingly, unsupervised svaseq has lower specificity (high false positive rate), empirical RUV, and with no adjustment has lower sensitivity (high false negative rate). We also observed similar trends when comparing ***BalanceIT*** library with the randomization library (**Figure S4A-B, c-d**), in which the ***BalanceIT*** library shows better DEG performance, due to fewer false positive DEGs. Unsupervised svaseq has the overall enhanced performance due to the decreased false positive DEGs in the ***BalanceIT*** library. However, the low sensitivity (high false negative) of empirical RUV with no adjustment did considerably improve upon the ***BalanceIT*** library design. Studies have shown that batch effects induce widespread dependence in expression variation,[1,2] and, p-values of null genes would be skewed away from an uniform distribution in a significance analysis.[3] To investigate if the DEG performances could be attributed to the unwanted variation (clinical covariates) generated from library design, we further examined the p-value distribution of differential expression significance (**Figure S5)**. Apparently, a library without batches (known batch (orange)) shows uniformly distributed p-values, but a library with batches (no adjustment (yellow)) shows a skewed distribution. Importantly, the ***BalanceIT*** library shows the most uniform p-value distribution (***BalanceIT*** library > Balanced library > randomized library > imbalanced library) for the differential expression analysis of various batch effect adjusted gene expressions using PCA (blue), supervised/unsupervised svaseq (pink), and empirical RUV (green). These results suggest that a better-balanced library is more amenable to removing unwanted variation, resulting more accurate DEG analysis.

Lastly, we used the identified DEGs from different designs corrected by different batch effect correction methods to assess their performance in the downstream gene set enrichment analysis. Regarding the GSEA performance, we assessed the performance by *F*-score, which is the harmonic mean of precision and sensitivity (**Methods**). We found that the balanced library outperforms the imbalanced library, and the fully balanced ***BalanceIT*** library has better performance than the randomization library (**Figure S4C**). Thus, a better-balanced library design also yields more accurate enriched gene sets, while a less balanced library design may lead to erroneous conclusions from incorrect enriched gene sets.

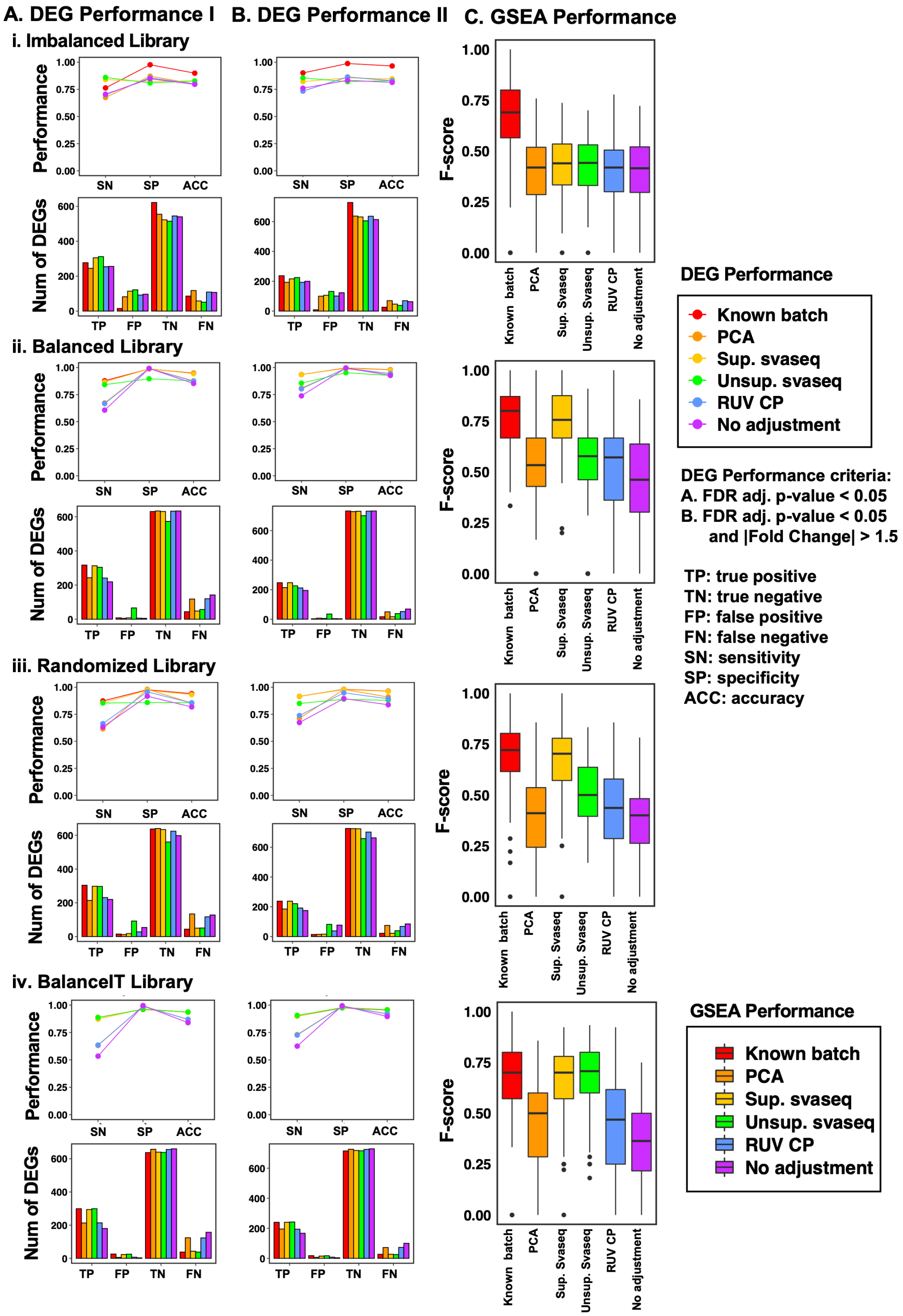


***Figure S4. Impact of different library designs on post-sequencing analysis.*** *The post-sequencing analysis are:* ***A)*** *differentially expression (DE) analysis with the DE criteria– FDR adjusted p-value < 0.05,* ***B)*** *differentially expression analysis with the DE criteria– FDR adjusted p-value < 0.05 and |Fold change| > 1.5, and* ***C)*** *gene set enrichment analysis. Here, we assessed the performance of post-sequencing analysis for the four different RNA-Seq library designs (see* ***Methods****):* ***i)*** *imbalanced library,* ***ii)*** *balanced library,* ***iii)*** *randomized library, and* ***iv)*** *BalanceIT library designs. Known batch (red), principal component analysis (PCA) (orange), supervised svaseq (yellow), unsupervised svaseq (green), RUV-CP: remove unwanted variation (RUV) using control probes (blue), and no adjustment (purple).*

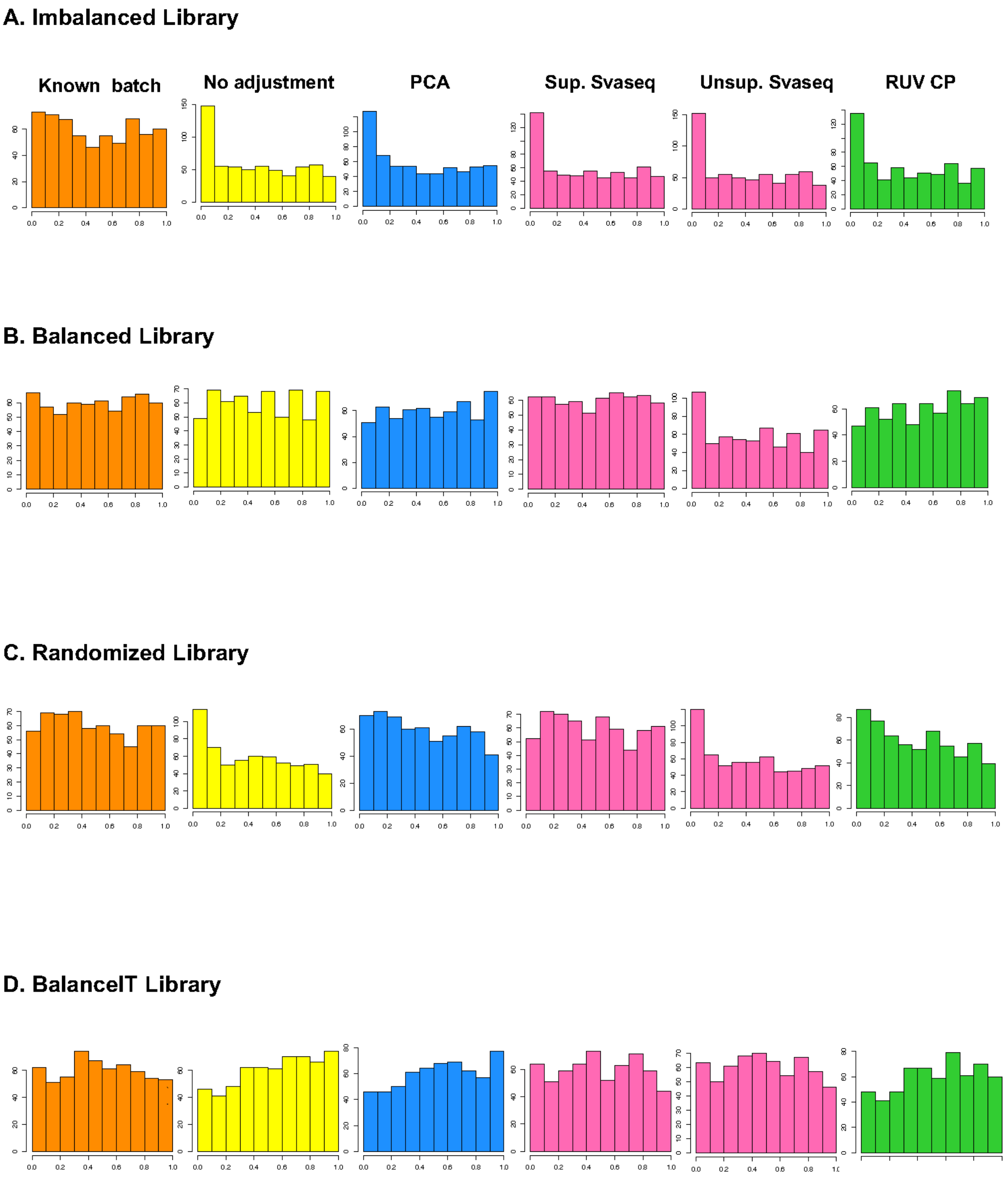


***Figure S5. P-value distribution for differential expression test between diagnosis groups (ASD vs. TD).*** *Four different strategies for the RNA-Seq data analysis:* ***A****) imbalanced library design,* ***B****) balanced library design,* ***C****) randomized library design, and* ***D****) BalanceIT library design. For each library design, the confounded gene expression was corrected by various batch effect correction methods: known batch (orange), no adjustment (yellow), principal component analysis (PCA) (blue), supervised svaseq (pink), unsupervised svaseq (pink), and remove unwanted variation (RUV) using control probes (green).*

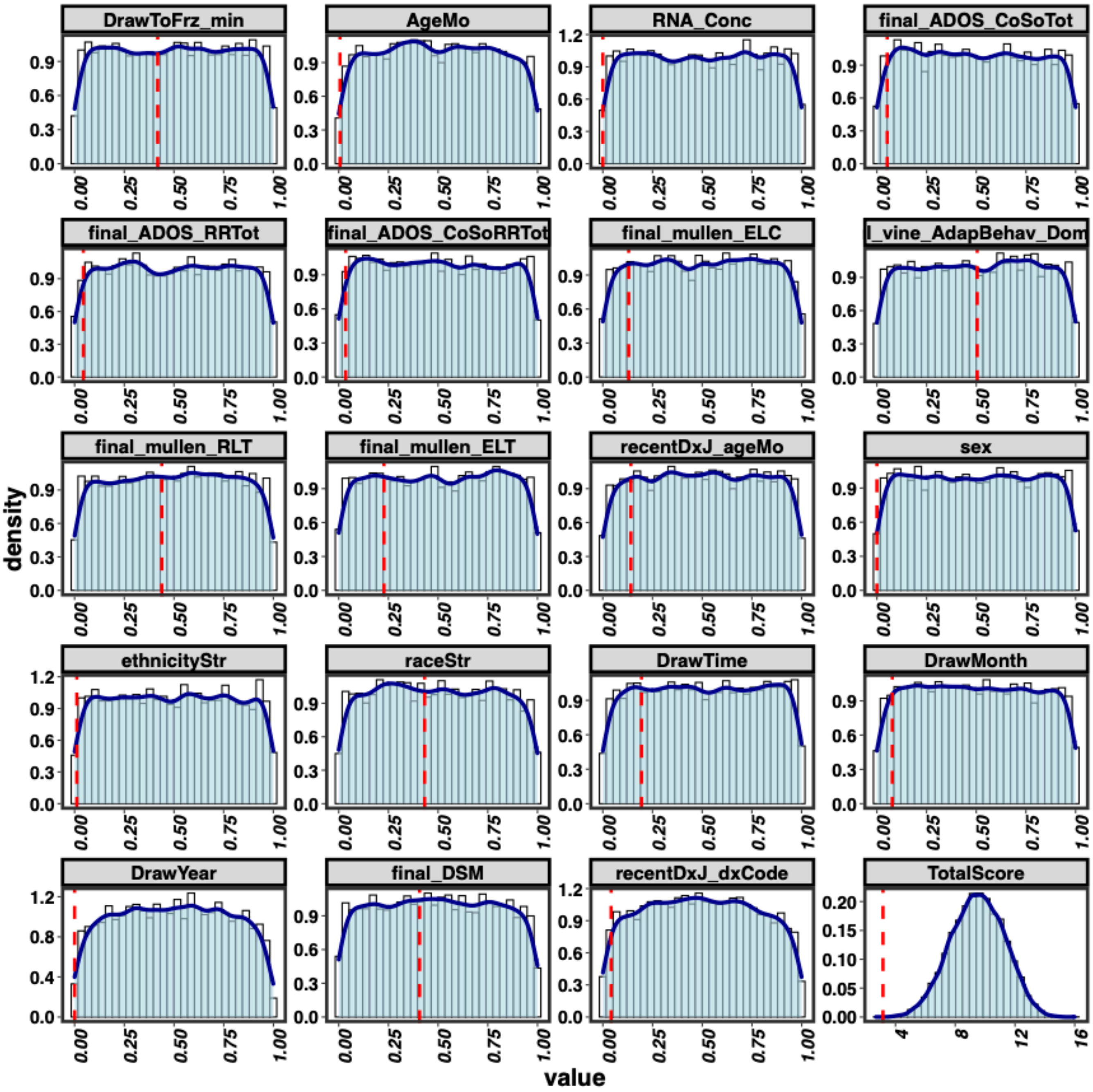


***Figure S6. Randomization test of the IBL library design for the ASD dataset with 678 samples.*** *We randomly assign the samples into different plates for 10000 times and assess the balance score for each covariate factor and total balance score for each new assigned library design. The red dashed lines indicate the balance score for the original IBL library design. The total score shows the original IBL library design is lowly balanced (total balance score=3.16 (normalized balance score=0.166;* ***Figure 3A****), p<0.001).*

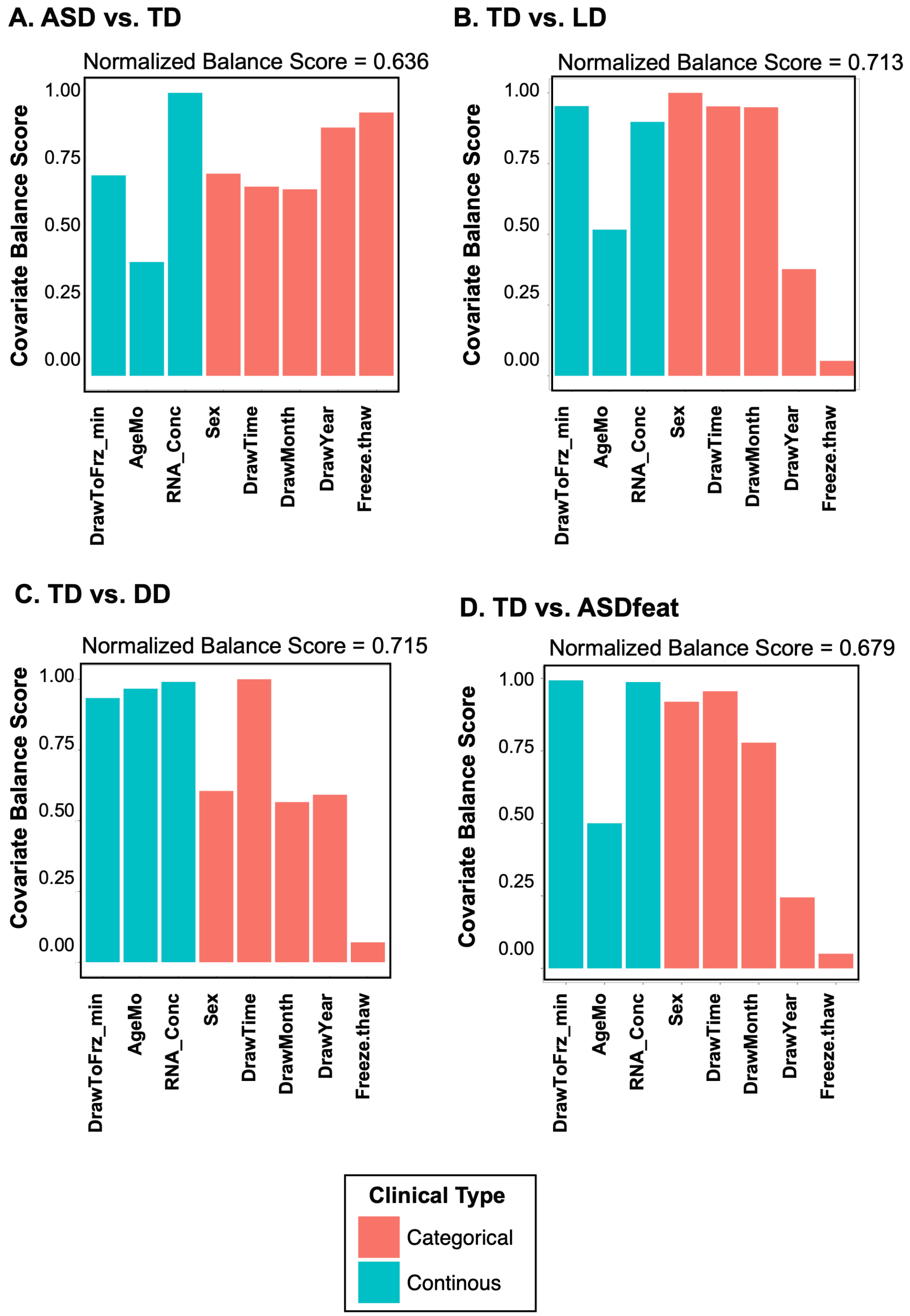


***Figure S7. Balancing results of the BalanceIT library design for the ASD dataset with selected 789 subjects.*** *The four between group balancing results are:* ***A****) ASD and TD,* ***B****) TD and LD,* ***C****) TD and DD, and* ***D****) TD and ASDfeat. The red bars denote categorical factors, and the turquoise bars denote continuous factors. The y-axis represents the balancing score of each factor. 'ASD': autism spectrum disorder, 'TD': typical developing, 'LD': language developmental delay, 'DD': developmental delay, and 'ASDfeat': ASD feature subjects.*

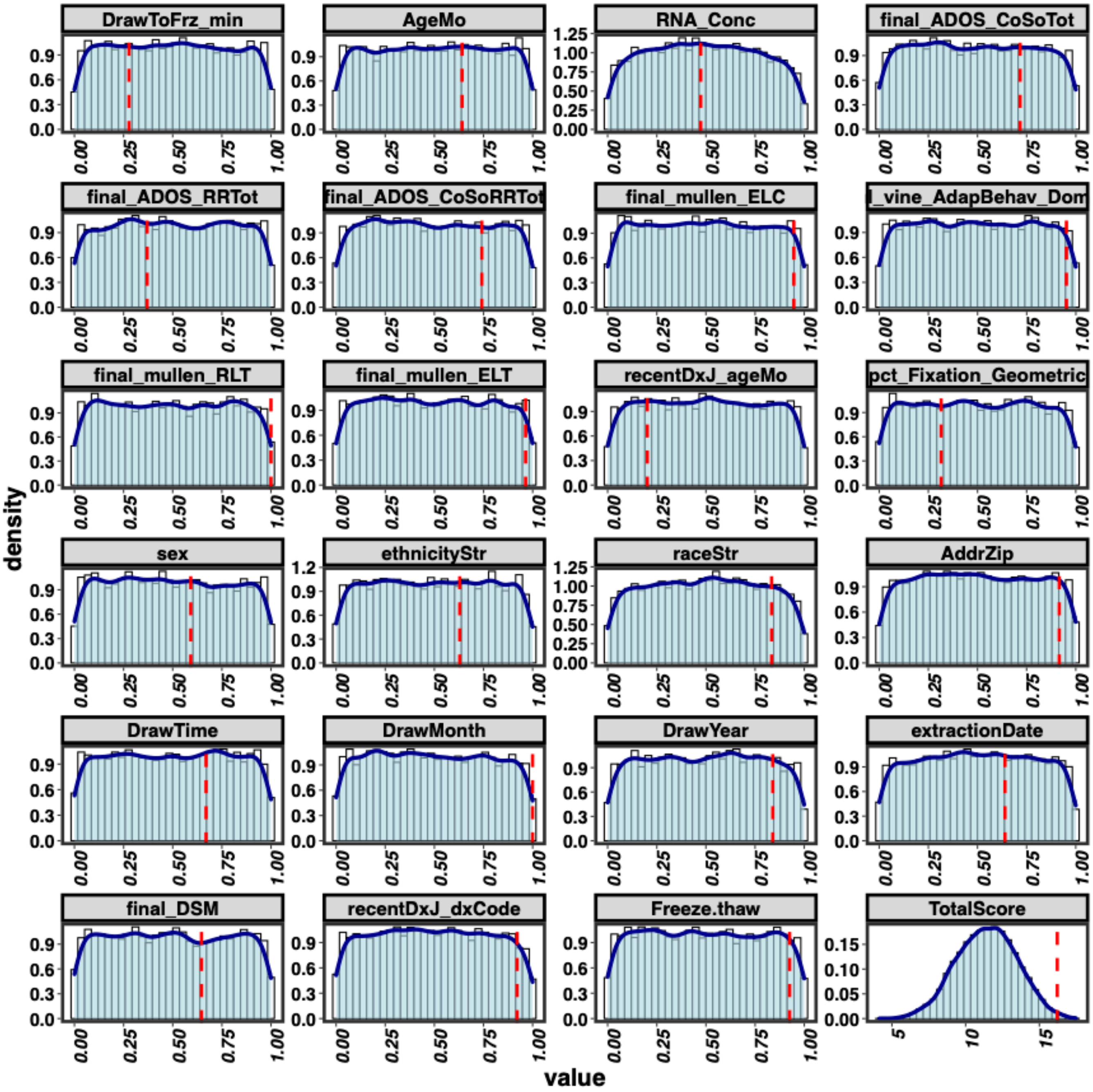


***Figure S8. Randomization test of the ACE-MRI library design for the ASD dataset with 358 samples.*** *We randomly assign the samples into different plates for 10000 times and assess the balance score for each covariate factor and total balance score for each new assigned library design. The red dashed lines indicate the balance score for the original ACE-MRI library design. The total score shows the original ACE-MRI library design is lowly balanced (total balance score=16.21 (normalized balance score=0.694;* ***Figure 4B****), p value=6.6e-03).*

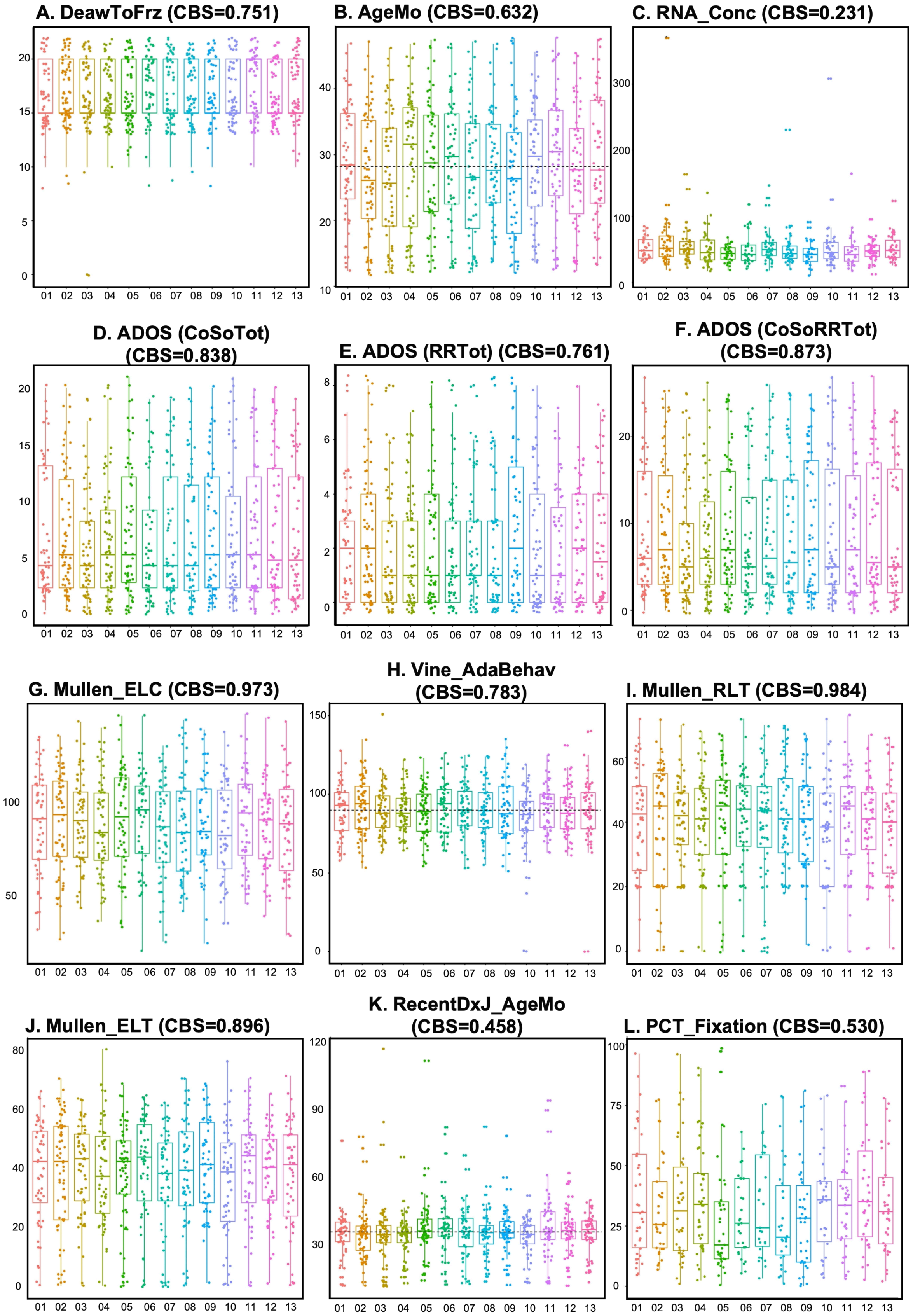


***Figure S9. Balancing results for the continuous covariates of the ASD dataset.*** *The x-axis represents the 13 plates. CBS: covariate balance score. The MRI-enriched plates are: plates #1, #4, #5, #6, #9,and #11.*

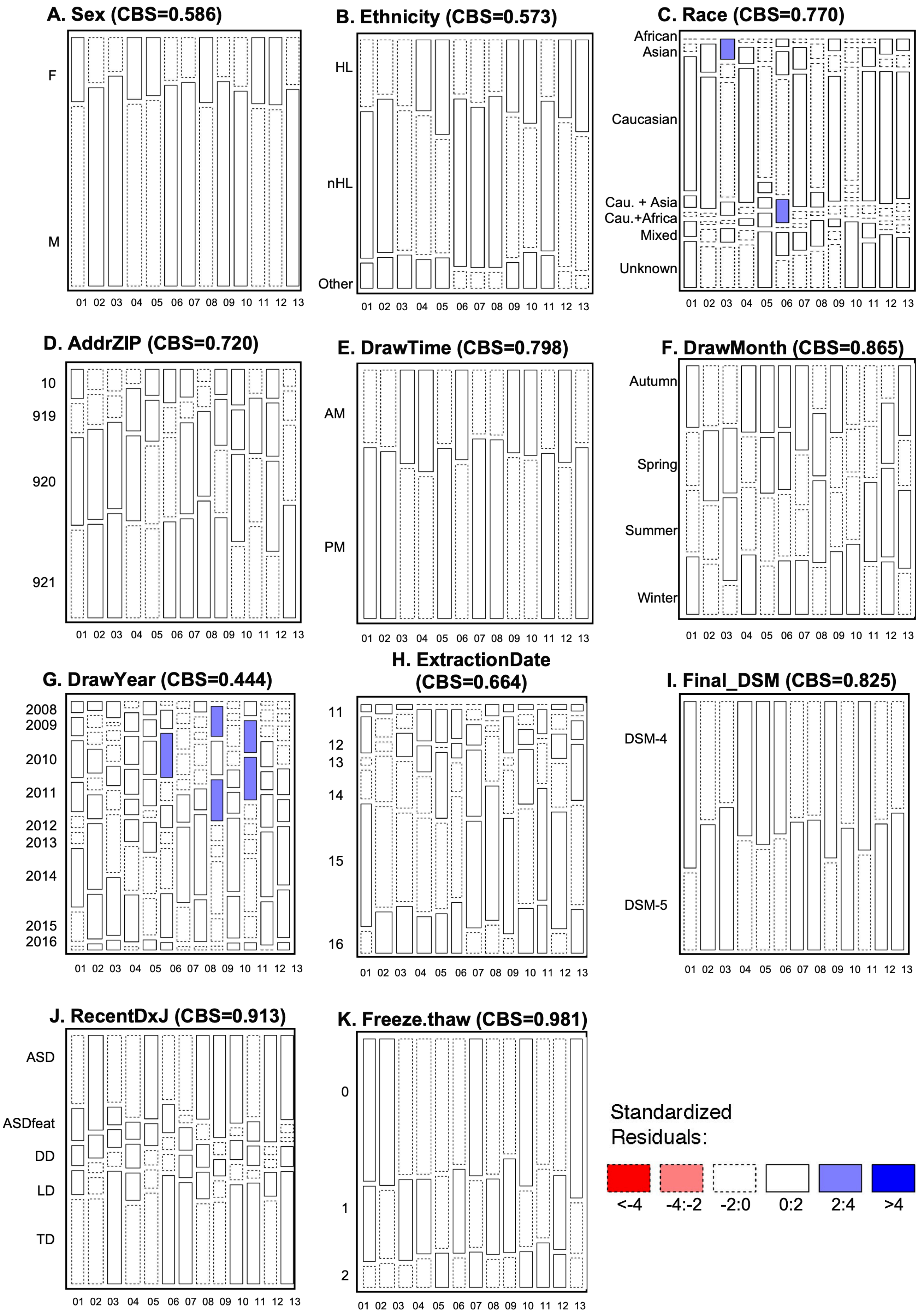


***Figure S10. Balancing results for the continuous covariates of the ASD dataset.*** *The x-axis represents the 13 plates. CBS: covariate balance score. The MRI-enriched plates are: plates #1, #4, #5, #6, #9,and #11.*

**Text S3. Validation on differential expression patterns for the IBL and ACE-MRI datasets**

From the balanced *ACE-MRI* library (789 unique toddlers) together with 266 longitudinal samples and 248 technical replicated samples (**Figures S11**), we conducted RNA-Seq from 549 subjects (6 of the 13 designed plates) to assess the performance of techniques for removing batch effects and performing downstream analyses (**Table S1**). We started by validating the expression patterns of ASD and TD samples in two independent cohorts. Our *ACE* dataset was composed of 389 subjects (n= 197 ASD and 192 TD subjects), and the *IBL* dataset was composed of 349 subjects (n= 220 ASD and 129 TD subjects). We first compared the DEGs from the IBL and ACE datasets to two previously published microarray studies. The first cohort (*tData*) was reported by Pramparo et al.[4], which was composed of 128 subjects (n= 84 ASD and 44 TD subjects). The second cohort (*NatNeur253*) was reported by Gazestani et al.[5], which was composed of 253 subjects (n= 84 ASD and 44 TD subjects). We first confirmed the DE pattens of *NatNeur253* and *tData* are reproducible from previous studies[5] (**Figure S12**). Our results further showed that the fold change comparison between *ACE* and *tData* datasets demonstrated similar fold change patterns (Pearson's correlation coefficient: 0.74 (only the shared DEGs) and 0.63 (all the DEGs); the left panels of **Figure S13, A** and **B**). However, we observed a much lower conservation of fold change patterns the for the comparison between the *IBL* and *tData* datasets (Pearson's correlation coefficient: 0.43 for shared DEGs and 0.59 for all DEGs). Similar trends of fold change conservation were observed when comparing to *NatNeur253*, with better conservation for the more balanced *ACE* dataset (the left panels of **Figures S14, A** and **B)**. These lower correlations are expected due to differences in platform (RNA-Seq for IBL and ACE, microarray for tData and NatNeur253), and other differences in the cohorts (male and female in IBL and ACE, while only male in tData and Nat Neur253).

To assess whether the sequencing plate batch effect could be effectively handled, we first performed Combat[6] batch effect correction on the plate information of *ACE* and *IBL* datasets. The conservation of fold change patterns showed mild improvement (3%-5%) on both datasets (right panels of **Figures S13A** and **5B**). These results are consistent with the simulated gene expression data analyses (**Figures 1 and S1**) wherein the imbalanced library design gene expression cannot be accurately corrected by the batch effect correction methods. Next, we focused our comparisons only on the male subjects of *ACE* and *IBL* datasets. Interestingly, we observed increased fold change conservations for *IBL* datasets on the male-only subjects (Pearson's correlation coefficients: 0.77 (common DEGs) and 0.81 (all DEGs); **Figure S13, C** and **D**) in contrast to all subjects (Pearson's correlation coefficients: 0.46 (common) and 0.64 (all)); **Figure S13, A** and **B**). Importantly, we found fold change conservation of *ACE* data can be easily controlled using Combat (Pearson's correlation coefficient: 0.83 (common DEGs) and 0.74 (all DEGs); **Figure S13, C** and **D**). Moreover, since gender is the primary confounding factor of the IBL dataset, focusing on males in the IBL dataset made the dataset more balanced. Not surprisingly, its conservation did not further improve by the Combat correction. We subsequently compared the ACE and IBL data to a second microarray dataset (NatNeur253) and go qualitatively similar results (see **Figures S14, C** and **D**). Thus, these analyses corroborated the effectiveness of balanced library design on batch effect control in clinical RNA-Seq data.

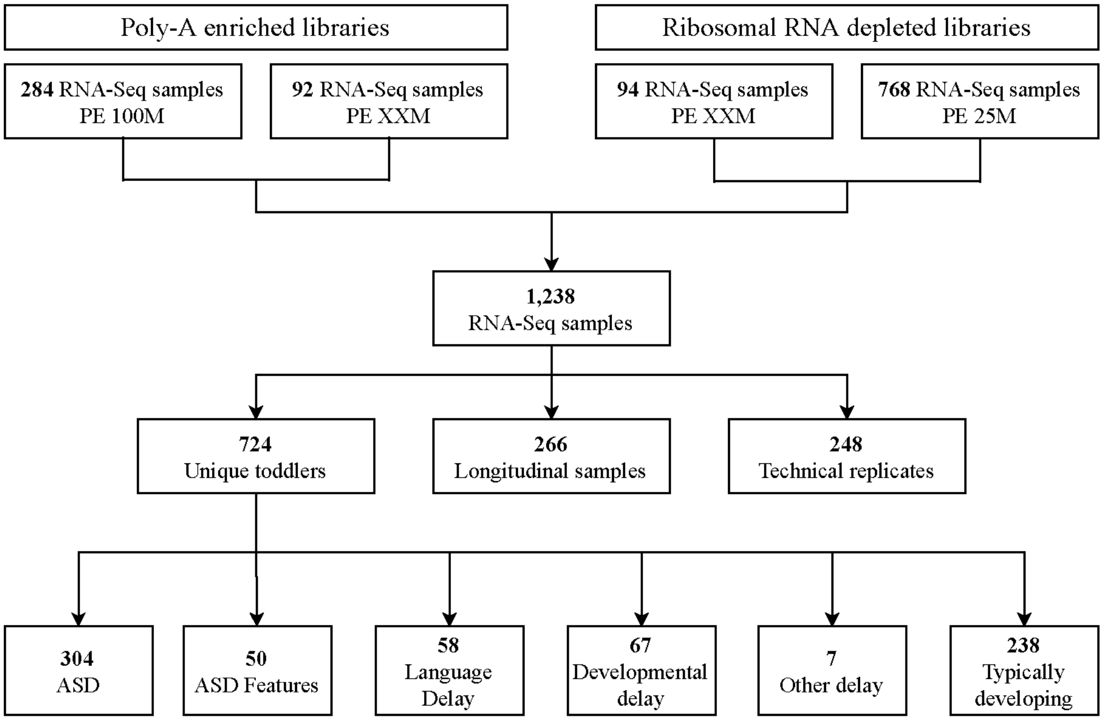


***Figure S11. Flow chart of the ACE balanced library in the RNA sequencing experiment.*** *In this study, we sequenced a total of 1,238 samples in 13 plates, including: 724 unique toddlers, 266 longitudinal samples, and 248 technical replicated samples.*

**Table S1. Overview of the 231 samples shared between the IBL dataset and the ACE-MRI dataset.**

| **Diagnosis** | | **Age in months** | |
|---|---|---|---|
| ASD | 145 | 14 Mo | 4 |
| TD | 86 | 15 Mo | 3 |
| **Sex** | | 16 Mo | 2 |
| Male | 157 | 17 Mo | 3 |
| Female | 74 | 18 Mo | 1 |
| **Batch** | | 19 Mo | 4 |
| p2321 | 37 | 20 Mo | 3 |
| p2327 | 36 | 21 Mo | 3 |
| p2341 | 26 | 23 Mo | 6 |
| p2350 | 75 | 24 Mo | 6 |
| p2354 | 32 | 25 Mo | 10 |
| p2395 | 25 | 26 Mo | 16 |
| **Year (Collection)** | | 27 Mo | 9 |
| 2008 | 16 | 28 Mo | 6 |
| 2009 | 36 | 29 Mo | 7 |
| 2010 | 48 | 30 Mo | 16 |
| 2011 | 67 | 31 Mo | 17 |
| 2012 | 55 | 32 Mo | 12 |
| 2013 | 9 | 33 Mo | 7 |
| **Percent rRNA** | | 34 Mo | 13 |
| 1 | 47 | 35 Mo | 10 |
| 2 | 150 | 36 Mo | 8 |
| 3 | 23 | 37 Mo | 7 |
| 4 | 1 | 38 Mo | 22 |
| 5 | 3 | 39 Mo | 9 |
| 6 | 2 | 40 Mo | 9 |
| 7 | 1 | 41 Mo | 5 |
| 8 | 1 | 42 Mo | 2 |
| 9 | 1 | 43 Mo | 2 |
| 10 | 1 | 44 Mo | 2 |
| 11 | 1 | 45 Mo | 2 |
| 12 | 0 | 46 Mo | 4 |
| 13 | 1 | 48 Mo | 1 |

A. NatNeur253 vs. DataC

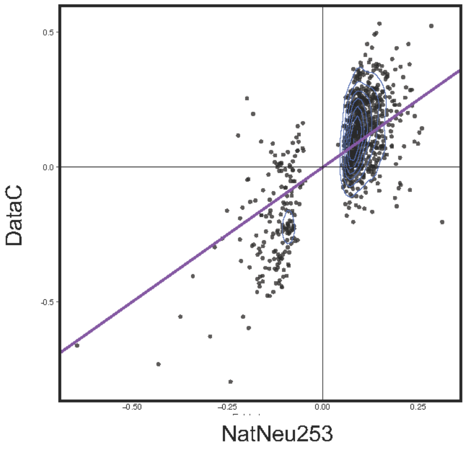


R=0.74

B. NatNeur253 vs. DataRep

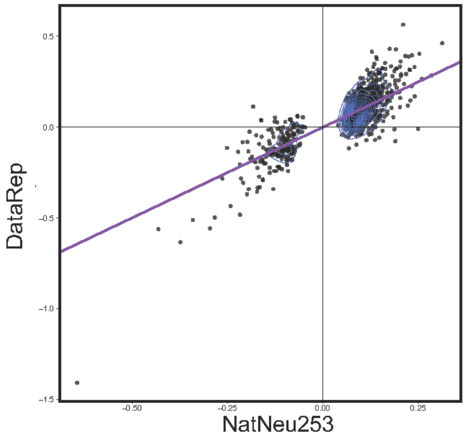


R=0.83

C. NatNeur253 vs. tData

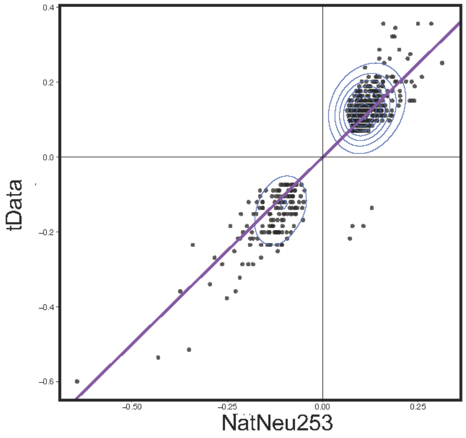


R=0.94

***Figure S12. Reproducibility analysis of observed DE pattens.*** *In this study, we reproduced the DE patterns from the published dataset:* ***A****) NatNeur253 and DataC,* ***B****) NatNeur253 and DataRep,* ***C****) NatNeur253 and tData.*

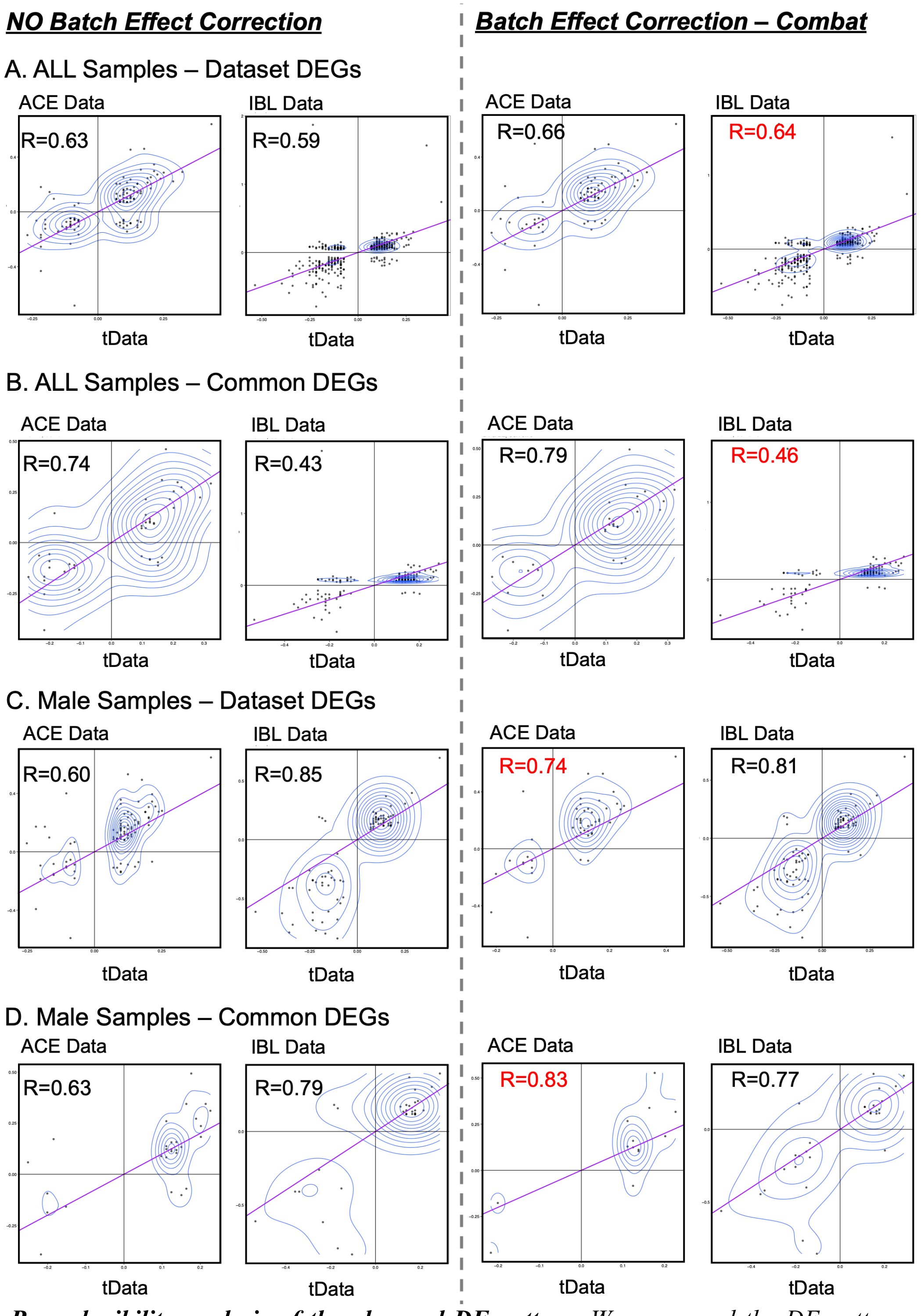


***Figure S13. Reproducibility analysis of the observed DE pattens.*** *We compared the DE patterns between our datasets (ACE and IBL) and the published tData dataset under different batch effect correction conditions (no correction (**left panels**) and Combat correction (**right panels**)). **A**) All DE genes of ACE/IBL data using all samples. **B**) Common DE genes of ACE/IBL data using all samples. **C**) All DE genes of ACE/IBL data using only the male samples. **D**) Common DE genes of ACE/IBL data using only the male samples.*

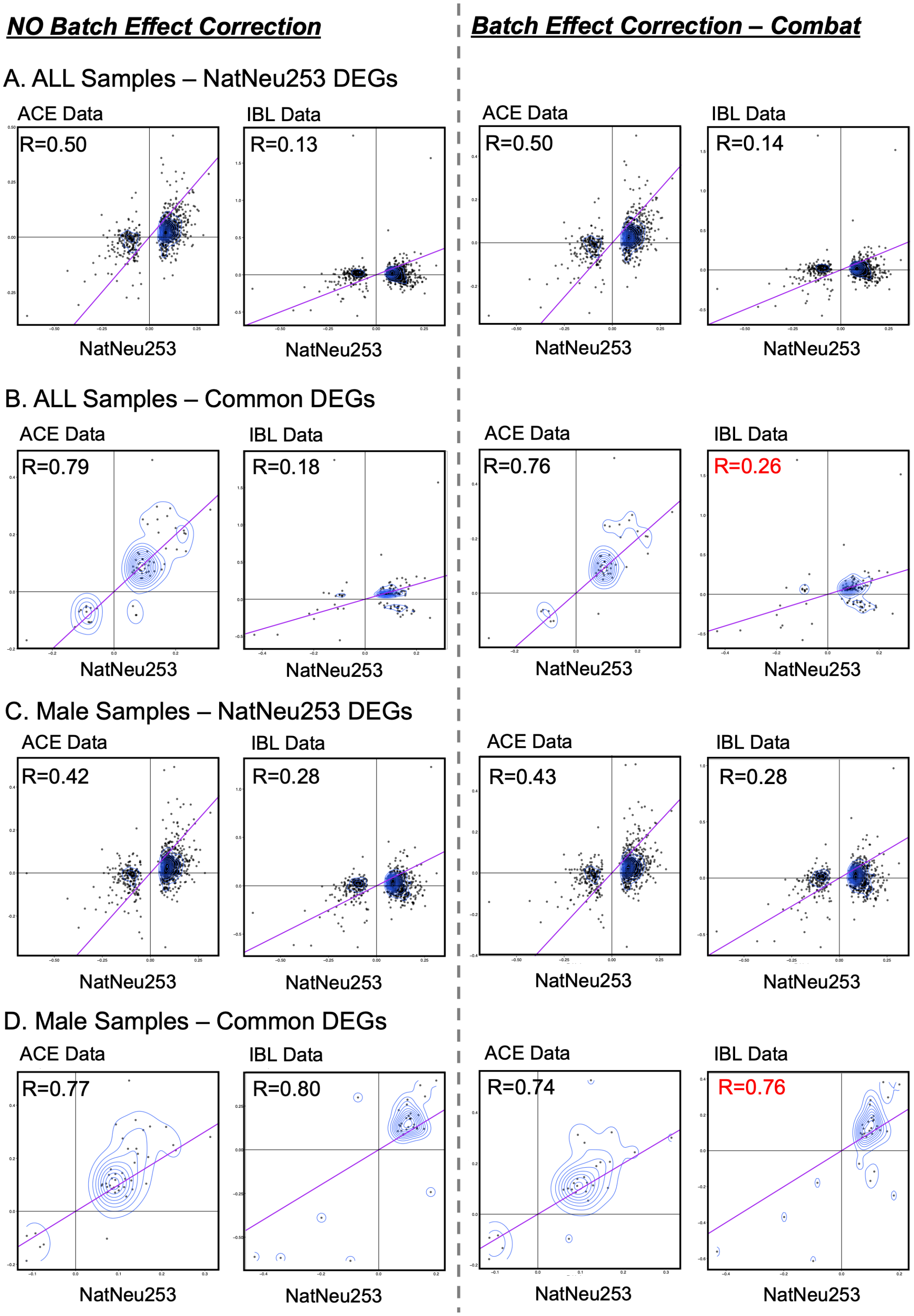


***Figure S14. Reproducibility analysis of the observed DE pattens.*** *We compared the DE patterns between our datasets (ACE and IBL) and the published NatNeu253 dataset under different batch effect correction conditions (no correction (**left panels**) and Combat correction (**right panels**)). **A)** All DE genes of ACE/IBL data using all samples. **B)** Common DE genes of ACE/IBL data using all samples. **C)** All DE genes of ACE/IBL data using only the male samples. **D)** Common DE genes of ACE/IBL data using only the male samples.*

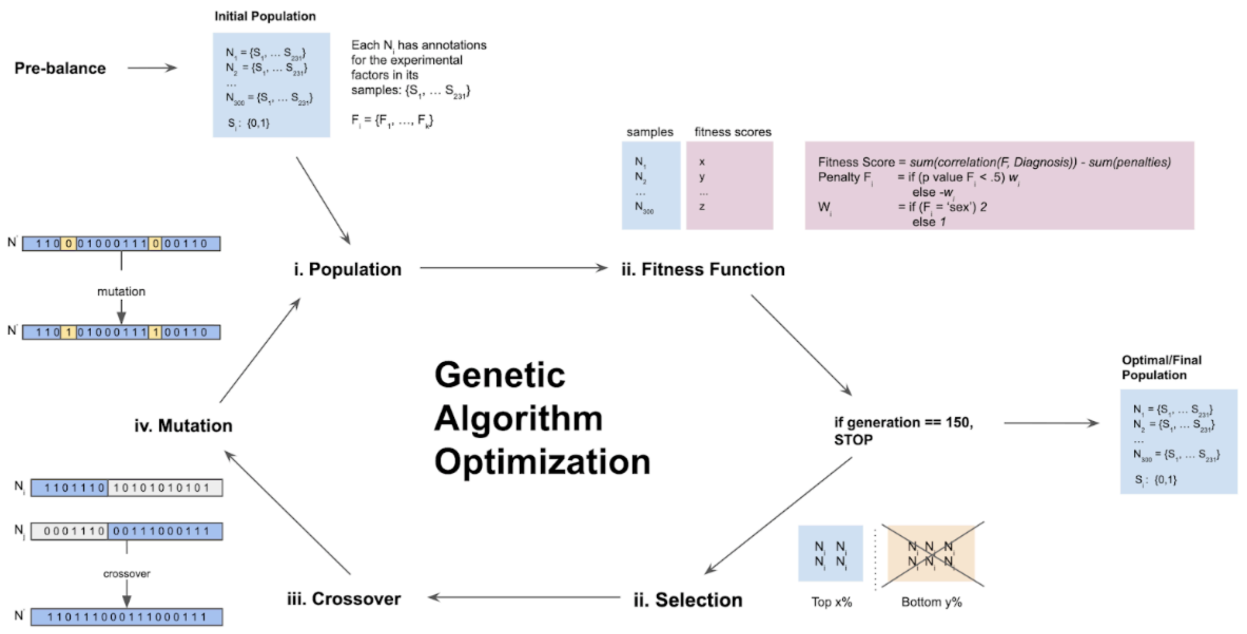


***Figure S13. Overview of the Genetic Algorithm Optimization framework of BalanceIT.*** *The GA-based optimization approach consists of six steps:* ***(i) initial population****–the initial population was generated by a set of 300 roughly pre-balanced solutions with sample size ranged between 150 to 165, termed as* $N=\{N_1, N_2, \ldots, N_{300}\}$*;* ***(ii) fitness function****–the function to assess how balance of the experimental factors in an individual solution* $N_i$*. For each* $N_i$*, each experimental factor is evaluated by performing the ANOVA test for continuous factors and by the Chi-square test for the categorical factors. The p-value of statistical test is considered as the 'balance score' of the testing experimental factor;* ***(iii) stop criteria****–the optimization process will be terminated after 150 generations;* ***(iv) selection****– top 30% of individuals served as candidates for further crossover and mutation to generate a new individual of the next-generation population;* ***(v) crossover****– the child individual is generated by swapping two parent chromosomes with a single crossover point; and* ***(vi) mutation****– a mutation occurs by randomly changing the value of a chromosome in the child individual to maintain population diversity.*
